# Spatiotemporal dynamics of ethylene biosynthesis shape infection and nodule initiation in Medicago truncatula

**DOI:** 10.64898/2026.03.06.710031

**Authors:** Sophia Müller, Thijs Stegmann, Kelvin Adema, Rens Holmer, Amber van Seters, Robin van Velzen, Olga Kulikova, Tristan Wijsman, Joel Klein, Josefina-Patricia Fernandez-Moreno, Anna N. Stepanova, Jose M. Alonso, Henk Franssen, Estibaliz Larrainzar, Arjan van Zeijl, Wouter Kohlen

## Abstract

Ethylene is a well-established negative regulator of nodulation, yet how ethylene biosynthesis and perception are spatially coordinated during early symbiotic signalling remains unresolved. Here, we investigate the dynamics of ethylene responses in *Medicago truncatula* using transcriptomics, promoter–reporter analyses, loss-of-function approaches and a synthetic reporter. We show that the activity of the ethylene-responsive *EBSn* reporter shifts from inner root tissues under non-symbiotic conditions to the outer cortex and epidermis following rhizobial inoculation, revealing a spatial reprogramming of ethylene signalling. Among the eight Medicago *1-AMINOCYCLOPROPANE-1-CARBOXYLIC ACID SYNTHASE (ACS)* genes, upon rhizobia application *MtACS3* is induced in outer root cell layers, while *MtACS10* is repressed in the inner cortex and pericycle, mirroring the shift in ethylene perception. Functional analysis demonstrates that MtACS10 restricts nodule initiation, whereas *MtACS3* modulates infection thread number, prevents nodule clustering, and contributes to radial positioning of nodule primordia. Rhizobial induced ectopic *ACS* expression in the root interior counteracts *MtACS10* repression and blocks nodulation, highlighting the requirement for spatially confined downregulation of ethylene biosynthesis. Together, these findings establish a framework in which localized shift in ethylene biosynthesis, mediated by distinct Medicago *ACS* genes, balances infection and organogenesis while co-defining the spatial limits of the root susceptible zone.

## INTRODUCTION

Plants constantly face environmental challenges such as fluctuating water, nutrients, light, temperature, humidity, and biotic stresses. To cope with these dynamic conditions, plants finely tune growth and development through phytohormones, which transmit signals within tissues and activate transcriptional programs (Evert and Eichhorn 2013). Among these adaptations, legumes have evolved a remarkable developmental innovation: the ability to establish nitrogen-fixing symbioses with rhizobia under nitrogen-limiting conditions, forming specialized root organs called nodules (Roy et al. 2020). Rhizobia activate nodule initiation only in a specific region of the early root differentiation zone. This region is therefore referred to as the root susceptible zone (Bhuvaneswari et al. 1980). These nodules provide a controlled environment for bacteria to convert atmospheric nitrogen into ammonium, supplying the plant with essential nutrients at the cost of high metabolic energy (Ferguson et al. 2019; Roy et al. 2020). Because of this energetic expense, nodule formation is tightly regulated by hormonal signals that integrate internal and external cues (Ferguson et al. 2019; Lin et al. 2020).

Nodule organogenesis is initiated upon perception of rhizobium-derived lipochitooligosaccharide (LCO) signals, called Nod factors, which activate key plant transcriptional regulators, including the master symbiotic regulator *NODULE INCEPTION (NIN)*, and trigger localized accumulation of cytokinins such as isopentenyl adenine (iP) and trans-zeatin (tZ) (Breakspear et al. 2014; Larrainzar et al. 2015; Limpens et al. 2015; van Zeijl et al. 2015b; Reid et al. 2017; Gühl et al. 2021). These signals collectively induce cell divisions in the root pericycle and cortex, generating a nodule primordium (Timmers et al. 1999; Xiao et al. 2014). Concurrently, LCO perception initiates a rhizobial infection process that begins in responsive root hairs. Root hair curling traps rhizobia in infection pockets, from which cell wall-bound infection threads guide bacteria to the developing nodule primordium. Inside the nodule, rhizobia are released and develop into symbiosomes: host membrane-bound compartments that facilitate nutrient exchange between plant and microbe (Roy et al. 2020; Adema and Kohlen 2024).

In the model legumes *Medicago truncatula* (Medicago) and *Lotus japonicus* (Lotus), ethylene-insensitive mutants (*sickle/Mtein2*) display hyper-infection and excessive numbers of, often clustered, nodules (Penmetsa and Cook 1997; Oldroyd et al. 2001; Penmetsa et al. 2008; Miyata et al. 2013), showing that ethylene is a key regulator in these processes. In contrast to auxins and cytokinins, ethylene acts as a negative regulator of nodule initiation (Penmetsa and Cook 1997; Gonzalez-Rizzo et al. 2006; Penmetsa et al. 2008; Guinel 2015; van Zeijl et al. 2015b; Xiao et al. 2025). Additionally, in Medicago, several *1-AMINOCYCLOPROPANE-1-CARBOXYLIC ACID SYNTHASE (ACS)* genes are transcriptionally induced following LCO perception (Larrainzar et al. 2015; van Zeijl et al. 2015b; Herrbach et al. 2017), and elevated ethylene production has been reported in alfalfa (*Medicago sativa*) and Lotus within hours of inoculation (Ligero et al. 1986, 1987; Reid et al. 2018).

Beyond nodulation, ethylene is widely recognized for roles in plant development and stress responses. These roles include promoting fruit ripening, root growth inhibition, shoot thickening, leaf senescence, and defence against pathogens (Verma et al. 2016; Dubois et al. 2018; Salvi et al. 2021). In *Arabidopsis thaliana* seedlings, ethylene induces the classical “triple response”; a phenotype characterized by inhibition of hypocotyl and root elongation, radial organ swelling, and exaggerated apical hook formation (Knight et al. 1910; Merchante and Stepanova 2017). The study of this response has been instrumental in identifying key components of the ethylene biosynthesis and signalling pathway (Binder 2020).

Ethylene biosynthesis is tightly regulated, possibly due to its rapid diffusion. The pathway begins with methionine, which is converted to S-adenosylmethionine (SAM) by SAM SYNTHASE. SAM is then converted into 1-aminocyclopropane-1-carboxylic acid (ACC) in the cytosol by ACC SYNTHASE (ACS), often considered the rate-limiting step of ethylene biosynthesis (Kende 1993). ACC is subsequently oxidized to ethylene by ACC OXIDASE (ACO), but it has been suggested that this step can also occur non-enzymatically via free oxygen (Depaepe and Van Der Straeten 2020; Pattyn et al. 2021). Additionally, ethylene production can be modulated by the catabolism of ACC to α-ketobutyrate via ACC DEAMINASE (ACD) activity (Plett et al. 2009).

Ethylene perception occurs at the endoplasmic reticulum (ER) membrane through a family of ethylene-specific receptors. In Arabidopsis these include ETHYLENE RECEPTOR1 (ETR1), ETR2, ETHYLENE RESPONSE SENSOR1 (ERS1), ERS2, and ETHYLENE INSENSITIVE4 (EIN4), which can form both homodimers and heterodimers (Gao et al. 2008). The ethylene-binding domain faces the ER lumen, while the cytosolic side interacts with protein complexes containing the negative regulator CONSTITUTIVE TRIPLE RESPONSE 1 (CTR1) (Gao et al. 2008; Shakeel et al. 2013). In the absence of ethylene, CTR1 phosphorylates the C-terminal end of the highly conserved ETHYLENE INSENSITIVE2 (EIN2), maintaining the pathway in an inactive state (Ju et al. 2012; Bowman et al. 2019). Ethylene binding triggers conformational changes that release CTR1, allowing EIN2 dephosphorylation, cleavage, and nuclear translocation of its C-terminal fragment. In the nucleus, EIN2-C activates transcription factors ETHYLENE INSENSITIVE3 (EIN3) and EIN3-LIKE1 (EIL1), which in turn regulate second-tier transcription factors, ETHYLENE RESPONSE FACTORs (ERFs), propagating transcriptional responses that mediate diverse ethylene-dependent phenotypes (Qiao et al. 2012; Wen et al. 2012; Ma and Dong 2021). Loss-of-function mutations in *EIN2* cause near-complete ethylene insensitivity, highlighting the central role of the EIN2 protein in the pathway (Guzmán and Ecker 1990).

The small and gaseous nature of ethylene poses major analytical challenges, complicating its detection and hindering detailed functional characterization. Unlike other key phytohormones such as auxin, abscisic acid, and gibberellic acid (Rizza et al. 2017; Herud-Sikimić et al. 2021; Rowe et al. 2023), ethylene currently lacks a dynamic *in vivo* biosensor. Moreover, ACC-based sensors do not directly measure ethylene activity and therefore cannot distinguish canonical ethylene signalling from ethylene-independent roles of ACC, which has been shown to act as a growth regulator in its own right (Van de Poel and Van Der Straeten 2014). Nonetheless, ethylene responses have been effectively monitored in Arabidopsis using *35S::EIN3-GFP* and *35S::EIL1-GFP* reporter lines to track ethylene-dependent stabilization of EIN3/EIL1 proteins (Yanagisawa et al. 2003; An et al. 2010; Ju et al. 2015), as well as the *EBS::GUS* transcriptional reporter to visualize EIN3-mediated transcriptional activation in specific tissues (Stepanova et al. 2007). The *EBS* reporter has since been refined and optimized in subsequent studies (Fernandez-Moreno et al. 2024, 2025). Attempts to apply previous versions of the *EBS* reporter in Medicago have been unsuccessful, and without functional ethylene sensors in this species, the spatiotemporal basis of ethylene-mediated repression of nodulation remains unresolved. The recently developed *EBSn* (*EIN3-Binding Site New*) reporter, which exhibits strong ethylene responsiveness in Arabidopsis and *Solanum lycopersicum* (Fernandez-Moreno et al. 2025), holds considerable promise for extending such analyses to Medicago and other legumes.

In Medicago, early transcriptional activation of *ACC synthases (ACS)* following rhizobial signal perception suggests that ethylene biosynthesis is dynamically reprogrammed during nodule organogenesis (Larrainzar et al. 2015; van Zeijl et al. 2015b). However, while ethylene is recognized as a negative regulator of nodulation, the mechanisms governing its spatiotemporal regulation in roots remain largely unknown. Here we show that early symbiotic signalling in Medicago is accompanied by a spatial reorganization of ethylene biosynthetic potential along the root. Specifically, the ACC synthases *MtACS3* and *MtACS10* exhibit antagonistic transcriptional regulation and distinct spatial expression domains following rhizobial perception. Genetic and physiological analyses indicate that these genes differentially modulate nodulation, influencing nodule number, positioning, and infection dynamics. In addition, our results support a role for ethylene signalling in defining the size and competence of the root susceptible zone. Together, these findings suggest that localized control of *ACS* gene expression contributes to the spatial coordination of nodulation.

## RESULTS

### The *EBSn* promoter is functional in the Medicago root and shows a spatial shift in activity during early symbiotic signalling

To investigate the spatial dynamics of ethylene signalling during early rhizobial interactions, we generated composite Medicago plants with transgenic roots expressing the synthetic ethylene-responsive reporter *EBSn::GUS* (Fernandez-Moreno et al. 2025). Composite plants have normal shoots and partially transgenic roots grown from a callus—used to study root-specific gene functions and phenotypes (Limpens et al. 2004). Under non-symbiotic (mock) conditions, strong *EBSn* promoter activity was observed in the inner tissues of the root susceptible zone (Fig. 1A, C). Upon inoculation with *Sinorhizobium meliloti* 2011 (Sm2011), *EBSn* activity shifted within 24 hours to the outer root layers (Fig. 1B). At 48 hours post inoculation (HPI), promoter activity was mainly restricted to the epidermis and developing nodule primordia (Fig. 1C).

**Figure 1.**
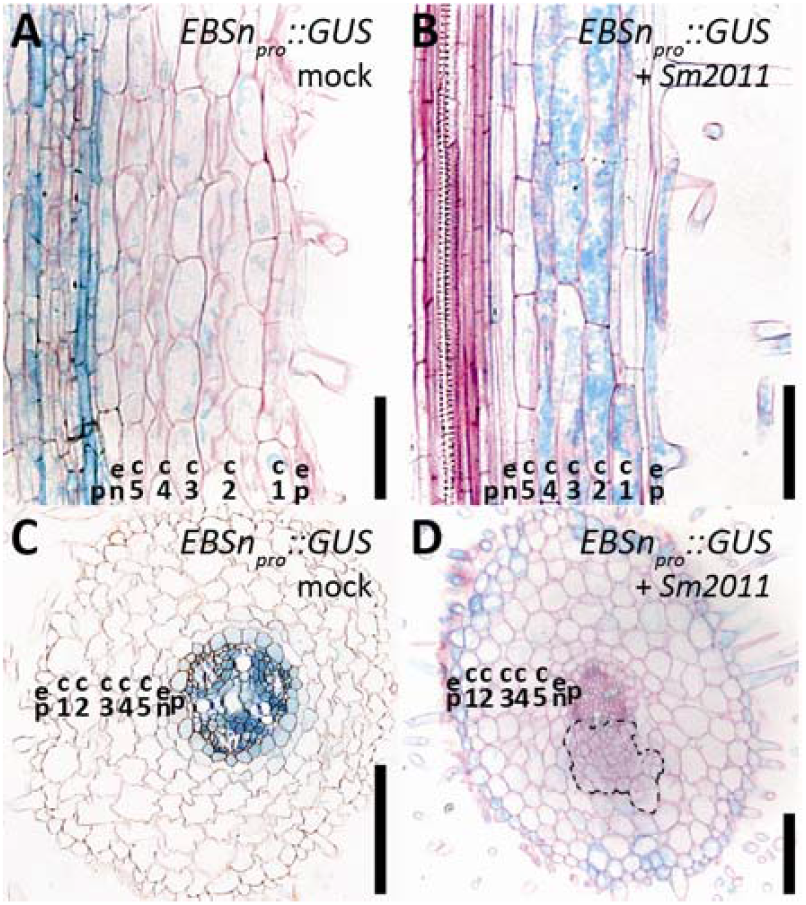
Ethylene responses in the root susceptible zone of Medicago during rhizobia-induced signaling. (A–C) Activity of the synthetic ethylene-responsive reporter *EBSn::GUS* (A) after 24 h mock treatment, (B) at 24 h post-inoculation (24hpi) with *Sm2011* (*Sinorhizobium meliloti* 2011), (C) in a root cross-section at 48 h mock treatment, and (D) in a root cross-section at 48 hpi. Dotted line indicates developing nodule primordia, ep, epidermis;c1-c5, cortical cell layers; en, endodermis; p, pericycle. Scale bars 100 µm.

Given that ethylene is a gaseous phytohormone assumed to be capable of diffusing across tissues, we hypothesized that this distinct spatiotemporal pattern of *EBSn* activity reflects either a direct diffusion pattern or a shift in localized ethylene biosynthesis. Ethylene is synthesized via the sequential action of ACS and ACO (Kende 1993) (Fig. 2A). Previous phylogenetic analysis of the Medicago genome revealed five *ACO*, one *ACD* and eight *ACS* genes (Fig. 2B-D, supplemental file 1, supplemental table S1, Gómez-Fernández et al. 2025). To explore candidate biosynthesis genes possibly involved in the observed ethylene dynamics, we performed RNA-seq analysis on the root susceptible zone (a 5 mm region starting 5 mm from the root tip) treated with either mock or LCOs. In pea (*Lathyrus oleraceus*, formerly *Pisum sativum*), PsACO expression has been reported to correlate with nodule initiation (Heidstra et al. 1997). For this reason, we first focused on the Medicago ACO genes. In our system (FA medium, 0.25 mM NO_3_^−^, no addition of aminoethoxyvinylglycine (AVG)), several *MtACO* genes exhibited robust expression, including *MtACO1* (∼20 TPM), *MtACO2* (∼160 TPM), *MtACO3* (∼390 TPM), and *MtACO4* (∼45 TPM), while *MtACO5* expression was below 1 TPM (Fig. 2E). In addition, *MtACD1* was expressed at approximately 50 TPM (Fig. 2F). Apart from a reduction in *MtACO4* expression, none of these genes were significantly affected by early LCO-induced signalling (Fig. 2E-F).

**Figure 2:**
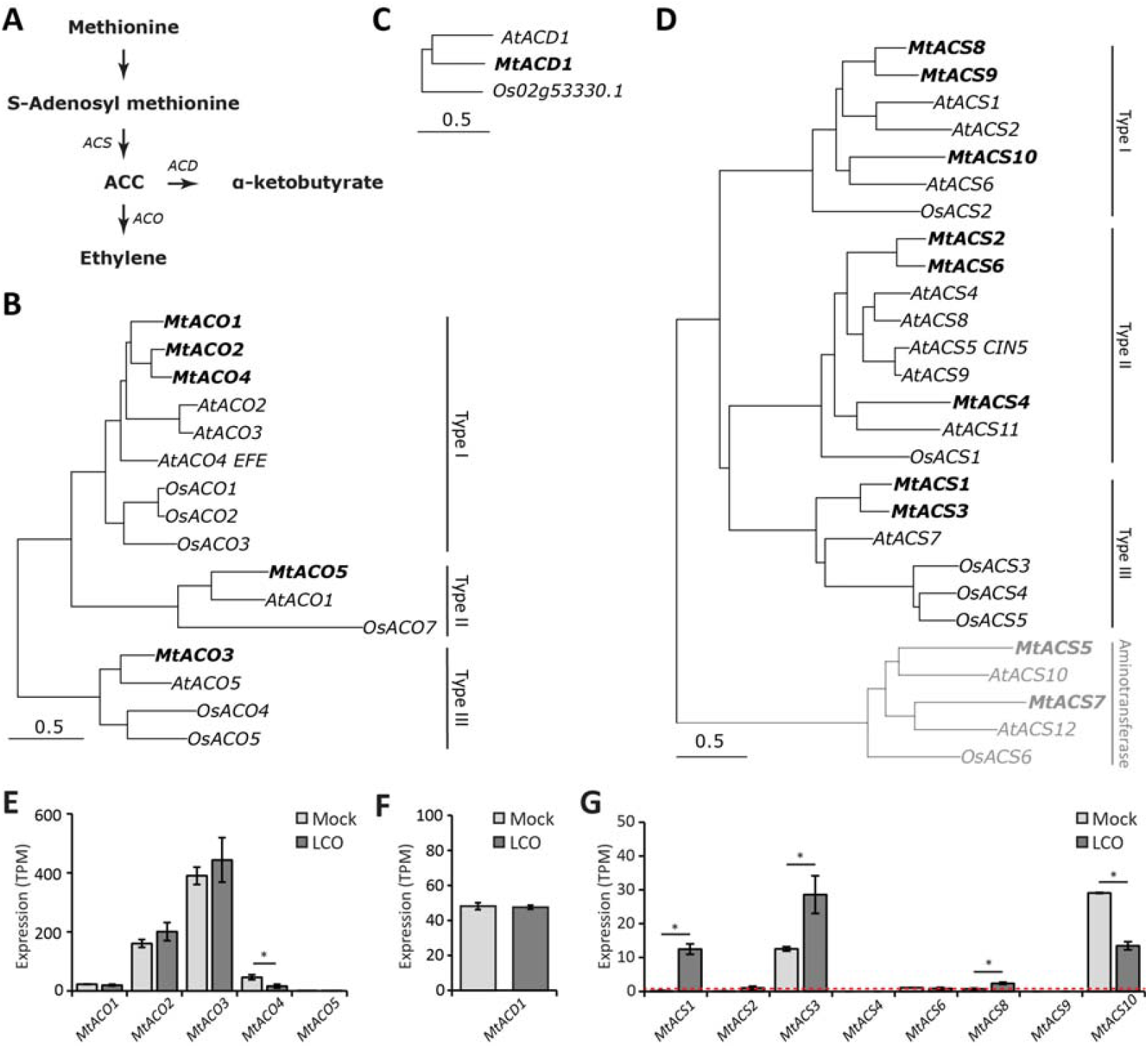
Identification and analysis of Ethylene biosynthesis genes in Medicago. (A) schematic representation of the ethylene biosynthesis pathway in plants, (B-D) Summary of a phylogenetic analysis of (B) ACC Oxidase (C) ACC deaminase and (D) ACC synthase amino acid sequences (*Mt = Medicago truncatula, At = Arabidopsis thaliana, Os = Oryza sativa*, for full phylogenetic trees including gene IDs see Supplemental file S2, in grey putative aminotransferases previously misidentified as ACC synthases (for review see Gómez-Fernández et al. 2025); scale bars indicate substitutions per site. (E-G) Expression of ethylene biosynthesis genes under control and lipo-chitooligosaccharide (LCO) treated conditions at 3 h post-LCO application. (E) *MtACO1, 2, 3, 4*, and *5;* (F) *MtACD*. (G) *MtACS1, 2, 3, 4, 6, 8, 9*, and *10*; Bars represent means ± SD (n = 2). Asterisks indicate significant differences between treatments (Student t-test, P < 0.05). Dotted line in (G) represents the used expression confident threshold (1 TPM, transcripts per million), data available at the European Nucleotide Archive (ENA) under accession number PRJEB38785; n, independent biological replicates each consisting of the susceptible zones of ∼16 pooled plate grown plants.

Next, we analysed the expression of the *ACS* genes. This revealed that under mock conditions, only *MtACS3* and *MtACS10* were expressed well above our expression confidence threshold of 1 TPM (12.5 and 29.1 TPM, respectively; Fig. 2G). Except for *MtACS6*, which was expressed at 1.1 TPM, all remaining *ACS* genes were expressed at levels well below 1 TPM (Fig. 2G). In contrast to the *MtACO* and *MtACD* genes, several of the *MtACSs* were differentially regulated by LCO signalling. Specifically, *MtACS1* (0.0 TPM to 12.2 TPM, mock vs LCO treatment respectively), *MtACS3* (12.5 to 28.2 TPM), and *MtACS8* (0.7 TPM to 3.1 TPM) were upregulated, whereas *MtACS10* was downregulated (29.1 to 13.4 TPM) (Fig. 2G, supplemental figure S1). These results are in line with changes in the expression previously observed for these genes (Supplemental figure S2, Larrainzar et al. 2015; van Zeijl et al. 2015b; Schiessl et al. 2019).

### *MtACS3* and *MtACS10* exhibit differential transcriptional regulation and spatial expression patterns in response to LCO signalling

We hypothesised that genes involved in the regulation of susceptibility need to be expressed prior to inoculation. *MtACS3* and *MtACS10* represent interesting candidates, as they were the only *ACS* genes expressed well above our confidence threshold in the *Medicago* root susceptible zone before nodulation and both responded transcriptionally to LCO signalling. Therefore, we focused our subsequent analyses on these two genes. An independent qRT-PCR experiment confirmed the antagonistic regulation of the *MtACS3* and *MtACS10* expression (Fig. 3A, B), a pattern that we also identified after re-analysis of a published dataset (Supplemental Figure 2A, Schiessl et al. 2019). This convergence across independent studies strengthens the robustness of the observation and indicates that antagonistic regulation of these genes is a consistent feature of the early nodulation response.

**Figure 3:**
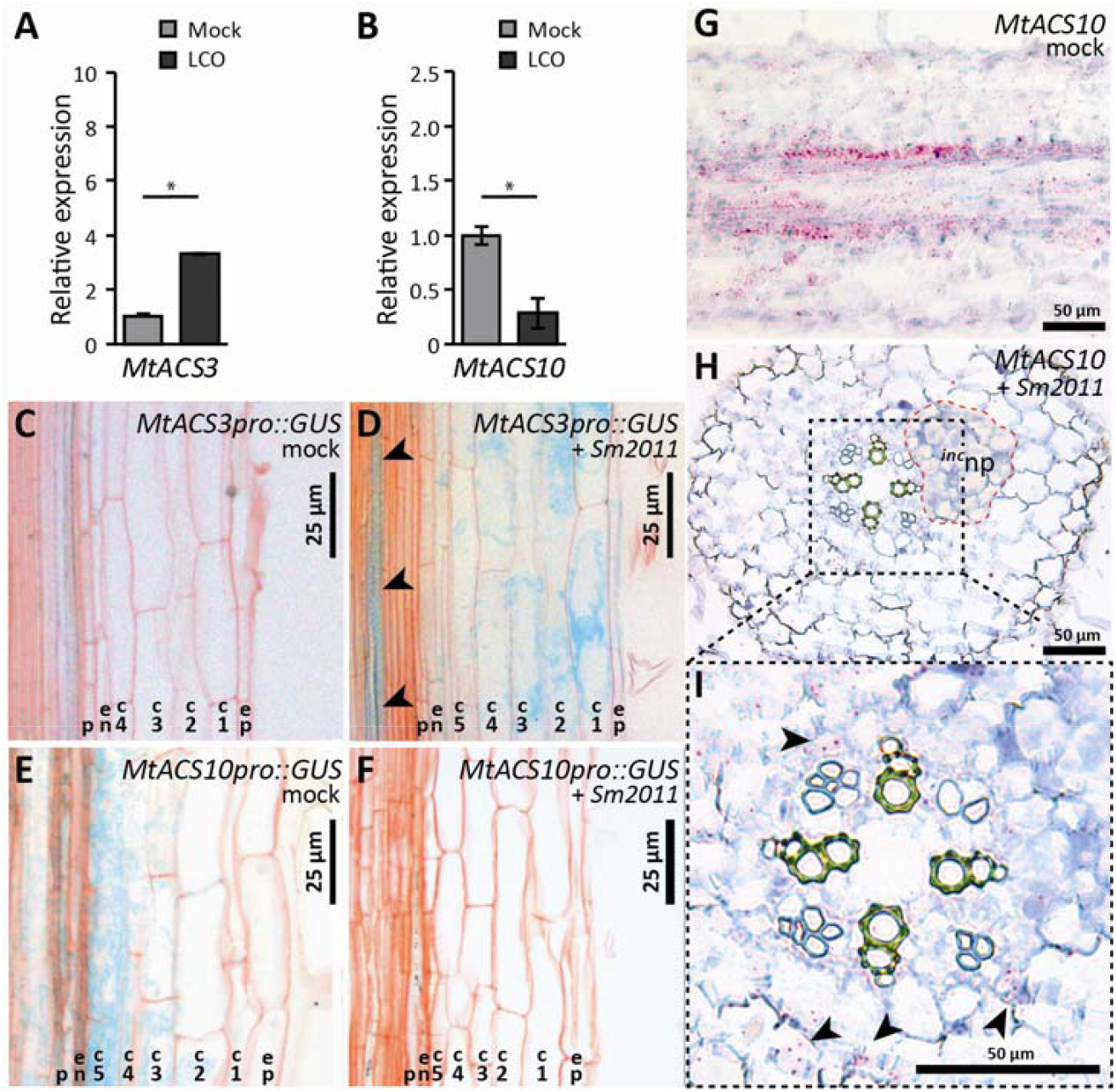
Expression Analysis of *MtACS3* and *MtACS10* in Medicago roots under symbiotic and control conditions. (A, B) Expression levels of (A) *MtACS3* and (B) *MtACS10* following mock and lipo-chitooligosaccharide (LCO) application, measured at 3 hours post-treatment. Bars represent mean ± SE (n=3, independent biological replicates each consisting of the susceptible zones of ∼16 pooled plate grown plants). Asterisks (*) indicate significant differences between treatments (Student t-test, *P* < 0.05). (C-F) *MtACS3* and *MtACS10* promoter activity and expression domains under mock and rhizobia inoculation. (C, D) *MtACS3pro::GUS* reporter activity at 20 hours post (C) mock or (D) *Sm2011* (*Sinorhizobium meliloti 2011*) spot-application, Arrows indicating GUS expression in the vasculature after (D) *Sm2011* treatment. (E, F) *MtACS10pro::GUS* activity at 20 hours post (E) mock or (F) *Sm2011* spot-application. (G, I) *MtACS10* expression domains in the pericycle and inner cortex as the result of RNA *in situ* hybridization. (G) Longitudinal section of an untreated Medicago wild-type (A17) root susceptible zone. (H-I) Cross-section of a root susceptible zone 36 hours post *Sm2011* application. (H) highlighting the position of an incipient nodule primordium (^*inc*^np) highlighted by the dotted red line, (I) Zoom in of vasculature and inner root cell layers. Hybridization signals are absent in reactivated pericycle and cortical cells of the incipient primordium. Hybridization signals appear as red dots (arrowheads in I), ep, epidermis;c1-c5, cortical cell layers; en, endodermis; p, pericycle in C-F).

Moreover, we found that the cumulative expression of *MtACS3* and *MtACS10* was similar between LCO-and mock-treated samples (Supplemental Figure S3A). This correlated with the observation that no significant differences in ACC levels could be detected in bulk samples from the root susceptible zone 3 hours after LCO application (Supplemental Figure S3B). These findings suggest that differential regulation of *MtACS3* and *MtACS10* can only influence nodule initiation if these two genes are expressed in distinct temporospatial domains.

To investigate this, we first determined the spatial expression patterns of *MtACS3* and *MtACS10*. To this end, we created composite plants bearing transgenic roots in which expression of the GUS reporter was driven by the putative promoter of either *MtACS3* or *MtACS10*. Analysis of the root susceptible zone revealed that, under mock conditions, the *MtACS3*_*pro*_*::GUS* reporter construct is mostly inactive in the root susceptible zone (Fig. 3C). As the first cell divisions have been reported to occur at 24 HPI (Xiao et al. 2014), we decided to analyse the effect of rhizobia spot inoculation just prior to this event. At approximately 20 HPI, the *MtACS3*_*pro*_*::GUS* promoter construct is activated in the vasculature, outer cortical cells, and the epidermis (Fig. 3D). In addition, the *MtACS10*_*pro*_*::GUS* reporter construct is active in the pericycle, endodermis, and inner cortical cell layers under non-inoculated conditions (Fig. 3E). When treated with Sm2011, the activity of the *MtACS10*_*pro*_*::GUS* reporter was abolished in all interior root cells close to the site of application of the root susceptible zone (Fig. 3F). Taken together, these reporter line observations suggest a spatial shift in the domain of ACS expression, and possibly ethylene biosynthesis.

To further investigate the cellular localization of *MtACS3* and *MtACS10* transcripts, we performed RNA *in situ* hybridization. RNA *in situ* hybridization against *MtACS3* revealed no signal in the root susceptible zone, although a *MtACS3* hybridization signal was detected in the epidermis and outer cortex at the elongation zone and start of the root differentiation zone (Supplemental Figure S4). This suggests that the *MtACS3* expression detected prior to LCO application in bulk samples likely reflects inclusion of adjacent tissue at the onset of the root differentiation zone, located just below the susceptible zone. Because the analysed root segments encompassed multiple developmental domains, transcripts originating from this proximal region were likely co-sampled. In contrast, *MtACS10* transcripts were predominantly detected in the pericycle and inner cortex of the elongation zone and the untreated susceptible zone (Fig. 3G, Supplemental Figure S4). Transverse sections of a 36 HPI *Sm2011*-treated root susceptible zone confirmed expression in a specific domain of inner cortical cells, surrounding the stele (Fig. 3H, I, arrowheads in I). However, *MtACS10* mRNA was excluded from the central vascular and incipient nodule primordia (Fig. 3H, I). Combined with the *MtACS10*_*pro*_*::GUS* reporter data, this suggests that upon inoculation with rhizobia, *MtACS10* is repressed in the cells that will initiate division to form a nodule primordia. Although we were unable to validate the observed promoter activity of *MtACS3* in the outer root layers by *in situ* hybridisation, results published by Breakspear at al., (2014) show *MtACS3* to be upregulated in the epidermis by both LCO and bacteria application in their root hair samples (Supplemental Figure 1B). Taken together, the observed *EBSn* pattern could thus, in part, be explained by *MtACS10* basal activity in the inner root tissues, followed by *MtACS3*, possibly in concert with other *MtACS* genes, induction in the outer cortex and epidermis after inoculation. This suggests that localized activity of these two ACS genes sequentially shapes the spatial dynamics of ethylene signalling during nodule initiation in Medicago roots.

### Different roles of MtACS3 and MtACS10 in the regulation of nodule development and infection thread formation in Medicago

Next, to investigate the functional role of MtACS3 and MtACS10 during nodulation, we generated composite plants with transgenic roots expressing RNA interference constructs targeting either *MtACS3 (ACS3*^*i*^) or *MtACS10 (ACS10*^*i*^). RNAi-mediated silencing specifically reduced *MtACS3* and *MtACS10* transcript levels by approximately 50% and 75%, respectively, without off-target effects on any of the other MtACS genes, nor the closely related putative aminotransferases (Supplemental Figure S5).

To assess whether knockdown of *MtACS3* or *MtACS10* was sufficient to alter nodulation, we inoculated these plants with Sm2011. At four weeks post inoculation (WPI), *ACS10*^*i*^ transgenic roots developed approximately twice as many nodules as either the empty vector control (EV) or the non-transgenic roots harvested from the same composite plants (Fig. 4A, Supplemental Figure S6). In contrast, *ACS3*^*i*^ did not exhibit a change in nodule number; however, transgenic roots in *ACS3*^*i*^ composite plants frequently formed fused nodules, a phenotype not often observed in EV nor *ACS10*^*i*^ roots (Fig. 4B). Given the transient nature of Agrobacterium rhizogenes–mediated root transformation and its limitations for quantitative analysis beyond microscopic phenotypes, we further investigated these traits using stable mutant lines.

**Figure 4:**
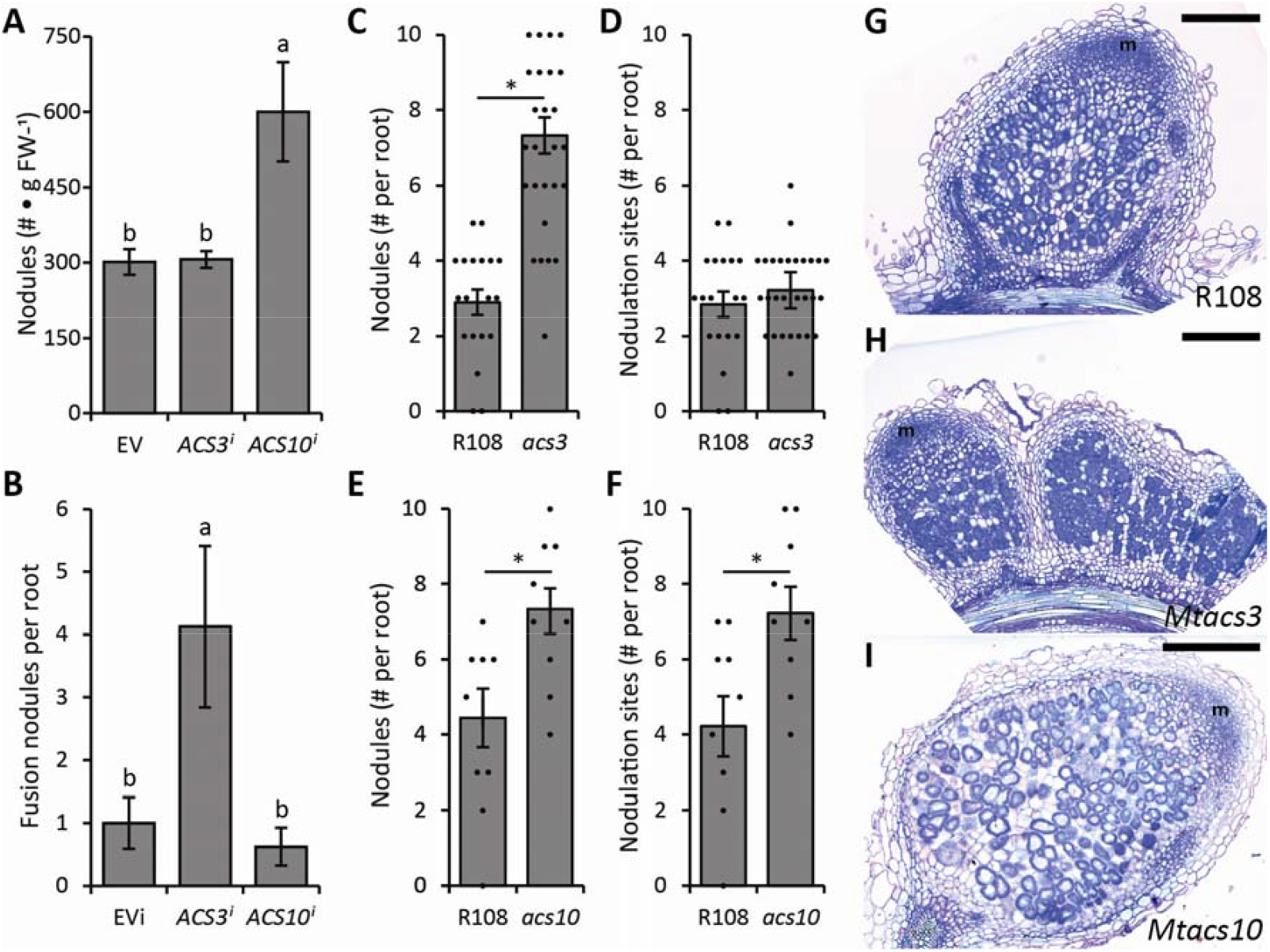
Nodulation phenotypes of *MtACS3* and *MtACS10* knockdown or knockout mutants. (A-B) Effect of *ACS3*^*i*^ or *ACS10* knockdown on nodulation in *Agrobacterium rhizogenes*-mediated composite plants grown in perlite (n>10, independently transformed root systems). (A) Average number of root nodules per gram FW. (B) Average number of nodule clusters per root system. Transformed and untransformed roots were separated based on *DsRed* expression (for untransformed roots, see Supplemental Fig. S5). Bars represent mean ± SE; different letters indicate significant differences (One-Way ANOVA followed by Tukey’s HSD test, *P* < 0.05). (C-F) Effect of *acs3* and *acs10* knockout mutations on nodulation in R108 plants grown on plates. (C) Average number of root nodules formed on R108 and *Mtacs3-1* (*acs3*) mutants (n>19, independent roots of plate grown plants). (D) Number of nodule initiation sites per root (n>19, independent roots of plate grown plants, same plants as in C). (E) Average number of root nodules formed on R108 and *Mtacs10-1* (*acs10*) mutants (n>8, independent roots of plate grown plants). (F) Number of nodule initiation sites per root (n>8, independent roots of plate grown plants, same plants as in E). Each dot represents an individual root, bars represent mean ± SE; an asterisk (*) indicate significant differences (Student t-test, *P* < 0.05). (G-I) Representative images of nodules formed on (G) wild-type R108, (H) *Mtacs3-1* mutant, or (I) *Mtacs10-1* mutant. Scale bars: 300 µm; m, nodule meristem.

We next screened the *Medicago Tnt1* retrotransposon insertion collection for mutants with disruptions in *MtACS3* or *MtACS10* (Tadege et al. 2008). We identified an insertion line in exon 2 of *MtACS3* and exon 1 of *MtACS10*, and designated them *Mtacs3-1* (NF0616), *Mtacs3-2* (NF1952), *Mtacs10-1* (NF15254), and *Mtacs10-2* (NF2329) (Supplemental Figure S7). When grown on plates, *Mtacs3-1* roots were significantly longer than those of wild-type R108, a phenotype reminiscent of the sickle root, and not observed in *Mtacs10-1* (Supplemental Figure S8A, B) (Penmetsa and Cook 1997).

To assess the functional contribution of MtACS3 and MtACS10 to ACC biosynthesis, we quantified ACC levels in distinct root zones. Given the differential expression patterns observed along the developmental root axis, roots were dissected into two segments: the root tip (RT; 0–5 mm from the apex) and the susceptible zone (SZ; 5–10 mm from the apex). In *Mtacs3-1* mutants, ACC levels in the root tip were reduced by approximately 75% relative to wild-type R108, whereas no significant reduction was observed in *Mtacs10-1* in this region (Supplemental Figure S9). Conversely, in the susceptible zone, ACC levels were reduced by ∼60% in *Mtacs10-1* compared to wild type, while *Mtacs3-1* showed no significant change (Supplemental Figure S9). These data indicate spatially distinct contributions of MtACS3 and MtACS10 to ACC biosynthesis along the root axis.

To evaluate nodulation in these mutants, *Mtacs3-1* and *Mtacs10-1* plants were inoculated with Sm2011 and analysed at 2 WPI. The *Mtacs3-1* mutant formed a substantial number of nodule clusters (Fig. 4C). Although total nodule number increased ∼2.5-fold in *Mtacs3-1*, the number of distinct nodulation sites per root was comparable to wild-type R108 (Fig. 4D). This observation is consistent with the *ACS3i* phenotype, which shows an increase in nodule clusters, and suggests a reduced capacity to regulate nodule number during initiation. This phenotype could reflect less constrained or excessive infection events, potentially contributing to the formation of closely spaced or clustered nodules. In agreement with the *ACS10*^*i*^ phenotype, *Mtacs10-1* mutant plants exhibited a ∼90% increase in nodule number compared to wild-type R108 (Fig. 4E), and, proportional to R108, only occasionally produced fused nodules when grown on plates. Both the number of nodules and the number of nodulation sites were similarly elevated in *Mtacs10-1* relative to wild type (Fig. 4E, F). Aside from nodule clusters in *Mtacs3-1*, nodules in both mutants appeared morphologically similar to wild-type R108 nodules (Fig. 4G–I). A similar effect on nodule numbers and clustering was observed for *Mtacs3-2* and *Mtacs10-2* mutants (Supplemental Figure S10).

Previous studies reported that the ethylene-insensitive *sickle* mutant forms a greater number of infection threads compared to wild-type plants (Penmetsa and Cook 1997). The role of ethylene during root hair infection was further supported by findings that treatment with the ethylene biosynthesis inhibitor AVG increased the number of infection threads per plant (Oldroyd et al. 2001). As we observed relatively high GUS activity in the epidermis at 48 HPI with Sm2011 during our EBSn reporter assays (Fig. 1), we hypothesized that the previously reported effects of ethylene on limiting surplus infections may be attributed to an increase in local ethylene biosynthesis in the outer cell layers of the root susceptible zone. To investigate whether *MtACS3* and/or *MtACS10* contribute to this response, we inoculated wild-type R108, *Mtacs3-1*, and *Mtacs10-1* lines with a Sm2011 strain constitutively expressing GFP, and quantified infection thread (IT) formation at two days post inoculation (DPI). On average, we detected two ITs per root in both R108, *Mtacs10-1* and *Mtacs10-2*. In contrast, both *Mtacs3-1* and *Mtacs3-2* roots exhibited approximately a fourfold increase in the number of ITs compared to wild-type R108 (Fig. 5, supplemental figure S11).

**Figure 5.**
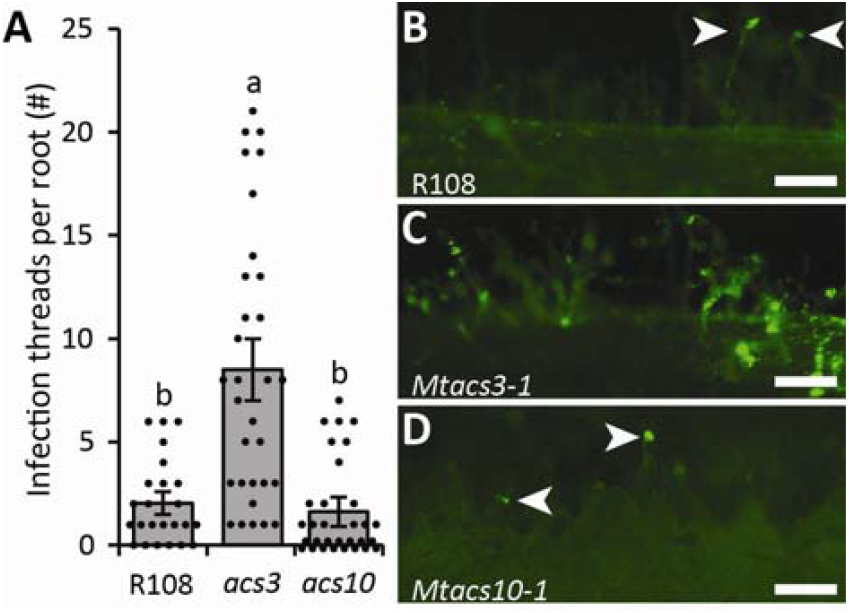
Effect of *Mtacs3-1* and *Mtacs10-1* loss-of-function mutations on infection thread formation. (A) Quantification of infection threads per spot-inoculation site following inoculation with GFP-labeled *Sm2011* (Sm2011-GFP) on wild-type R108, *Mtacs3-1* (*asc3*), *Mtacs10-1* (*acs10*). Each dot represents an individual spot inoculated susceptible zone (n>24); bars represent mean ± SE; different letters indicate statistical significance (one-way ANOVA followed by Tukey’s HSD test, *P* < 0.05). (B–D) Representative confocal images of Medicago (R108) roots at 2 days post-inoculation with *Sm2011*-GFP. (B) Wild-type R108, (C) *Mtacs3-1* mutant, and (D) *Mtacs10-1* mutant. Arrowheads indicate individual infection threads in (B) and (D); scale bars 150 µm.

In summary, *MtACS10* is expressed in the inner cortex pre-inoculation, and its mutant forms more nodules but a similar number of infection threads compared to wild-type R108. By contrast, *MtACS3* is induced in the outer root layers post-inoculation and likely functions in the outer cortex and epidermis, and its mutant shows both increased nodule numbers and elevated infection thread formation. Combined, the differential expression of *MtACS3* and *MtACS10* suggests an interesting, less straightforward, mechanistic link between infection threads and nodule primordium initiation that warrants further investigation.

### Loss off *MtACS3*, but not *MtACS10*, affects radial positioning of Medicago nodule primordia

Previous studies in pea reported that nodules form almost exclusively opposite the xylem poles, and that this spatial positioning is at least partially regulated by ethylene (Heidstra et al. 1997). A similar effect has been described for Medicago, though with slightly more variability: approximately 80% of nodules form opposite the xylem poles, while around 20% are located opposite the phloem poles (Penmetsa et al. 2003). Furthermore, the regulatory influence of ethylene appears to be more pronounced in Medicago, as the sickle mutant displays an approximately equal distribution of nodule primordia between xylem and phloem poles (Penmetsa et al. 2003).

To determine whether local ethylene biosynthesis by *MtACS3* or *MtACS10* contributes to radial positioning of nodules along the root axis, we grew wild-type R108, *Mtacs3-1, Mtacs3-2, Mtacs10-1* and *Mtacs10-2* lines in perlite inoculated with a Sm2011 strain constitutively expressing GFP. After 7 days, plants were harvested and the GFP signal was used to identify and isolate early rhizobia infection sites. This resulted in the collection of roughly 20–25 root segments per line containing early nodulation events. Root segments were fixed, embedded in plastic, and sectioned for analysis.

Although the central vasculature of Medicago roots can occasionally form a fourth xylem pole, all roots in this experiment developed the typical triarch configuration. In this arrangement, the three xylem poles are spaced approximately 120° apart, with a phloem pole located midway between each pair, at ∼60° intervals (Fig. 6). We sectioned all root segments and successfully traced 18–20 nodule primordia per genotype. Using ImageJ, we drew a line from the central vascular bundle through a xylem pole and a second line from the centre to the midpoint of the dividing cortical cells forming the primordium. The angle between these two lines was then calculated and categorized into six bins: 0– 10°, 10–20°, 20–30°, 30–40°, 40–50°, and 50–60° (Table 1). Primordia with angles <30° were classified as associated with the xylem pole, while those >30° were considered phloem associated.

**Table 1.** Angle distribution of the nodule primordia towards the xylem poles.

| Line | BIN1 (0-10°) | BIN2(10-20°) | BIN3 (20-30°) | BIN4 (30-40°) | BIN5 (40-50°) | BIN6 (50-60°) |
| --- | --- | --- | --- | --- | --- | --- |
| R108 | 13 | 5 | 2 | 0 | 0 | 0 |
| *Mtacs3-1* | 5 | 2 | 4 | 3 | 3 | 1 |
| *Mtacs3-2* | 3 | 3 | 2 | 5 | 0 | 2 |
| *Mtacs10-1* | 12 | 3 | 5 | 0 | 0 | 0 |
| *Mtacs10-2* | 10 | 2 | 2 | 0 | 0 | 0 |

**Figure 6:**
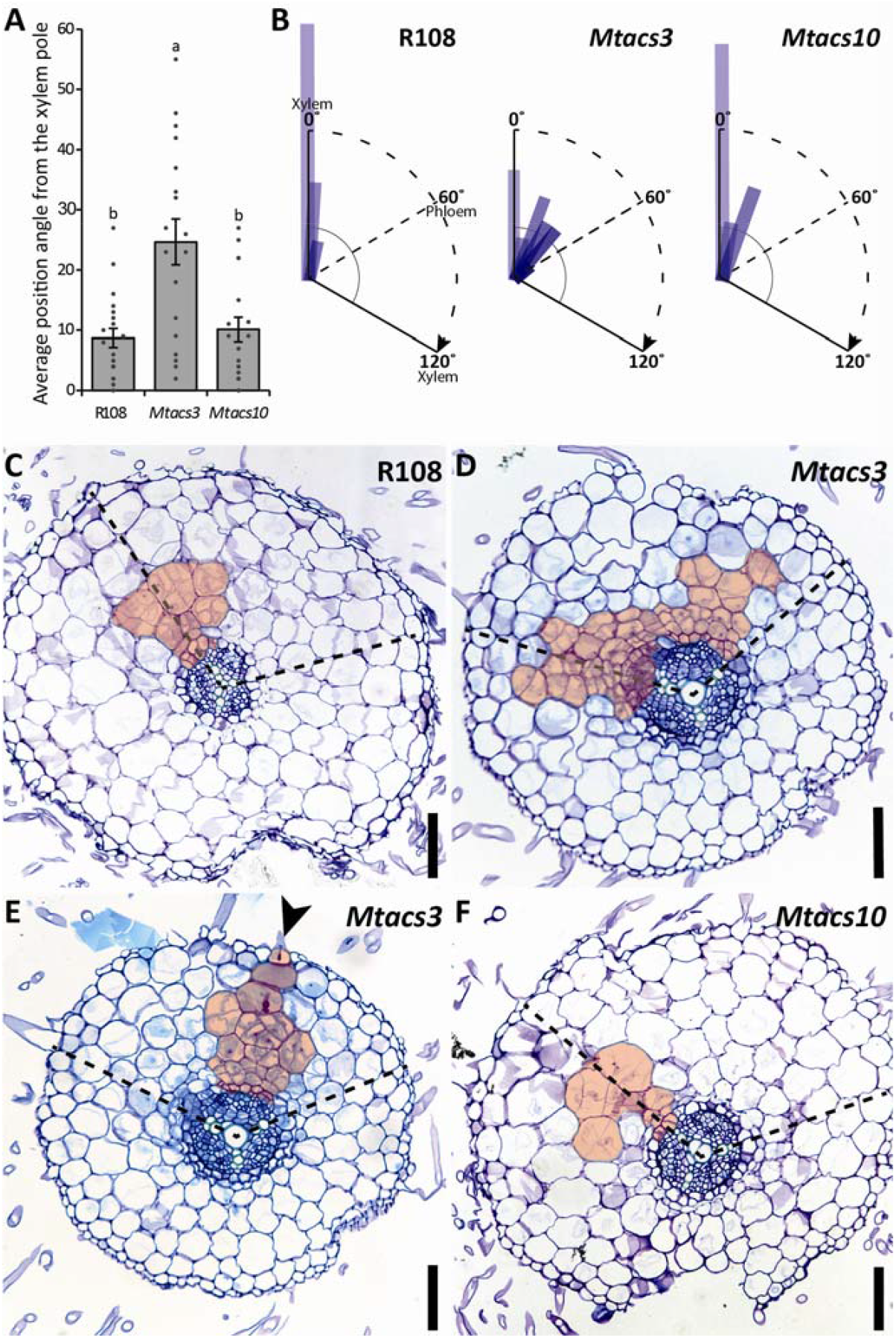
Nodule position in relation to xylem and phloem poles. (A) Average position angle of the nodule primordia from the xylem pole (B-D) schematic representation of one-third of the Medicago vasculature triarch, with a 120° angle between two xylem poles and the phloem in between at 60° showing (B) the relative positioning of nodule primordia at 3 days post-inoculation in wild-type R108, *Mtacs3-1*, and *Mtacs10-1* mutants. Each dot represents an individual sectioned susceptible zone (n>18); bars represent mean ± SE; different letters indicate statistical significance (one-way ANOVA followed by Tukey’s HSD test, *P* < 0.05). (C-F) Representative cross-sections of young nodule primordia (highlighted in red) 3 days after inoculation with a GFP-labeled *Sm2011* strain (*Sm2011-GFP*), formed on (C) wild-type *R108*, (D-E) *Mtacs3-1* and (F) *Mtacs10-1* mutants. Dotted lines indicate the two closest xylem poles. Arrowhead in E points at infection tread; scale bars 100 µm.

In wild-type R108, all nodule primordia were positioned opposite xylem poles, with the majority falling within the 0–10° range, averaging the angle between the xylem and nodule primordia at 8.7° (Fig. 6A-C, Table 1). In contrast, *Mtacs3-1* roots displayed a more randomized spatial distribution of nodule primordia, leading to an average angle of roughly 25° (Fig. 6A, B, D, E). In some cases, the primordium appeared to originate at a xylem pole but expanded across a broader radial domain (Fig. 6D), while in others, a clearly defined infection thread was observed initiating opposite the phloem pole (Fig. 6E, Arrowhead). *Mtacs10-1* roots, similar to R108, exhibited a strong preference for xylem pole-associated nodules, averaging the angle between the xylem and nodule primordia at 10.1° (Fig. 6A, B, F). Similar effects were observed for *Mtacs3-2* and *Mtacs10-2* (Table 1), demonstrating that the ethylene restriction on the radial positioning of nodules is co-depending on *MtACS3*, not *MtACS10*.

### Ectopic expression of *MtACS3* or *MtACS10* under control of the *MtCRE1* promoter leads to reduced nodulation

Next, we investigated whether the reduction in *MtACS10* expression could be linked to the root’s susceptibility to nodulation. To address this, we conducted a complementation experiment in which we inhibited the LCO-induced transcriptional repression of *MtACS10*. For this purpose, we leveraged the observation that *MtACS10* and *MtCRE1* are expressed in overlapping spatial domains (Fig. 3E, Gonzalez-Rizzo et al. 2006; Liu et al. 2019), and respond antagonistically to LCO application (Fig. 7A, B). While *MtACS10* transcript levels decrease by approximately 70% upon LCO treatment, MtCRE1 expression increases by roughly twofold (Fig. 7A), and their cumulative expression remains comparable between treatments (Fig. 7B).

**Figure 7:**
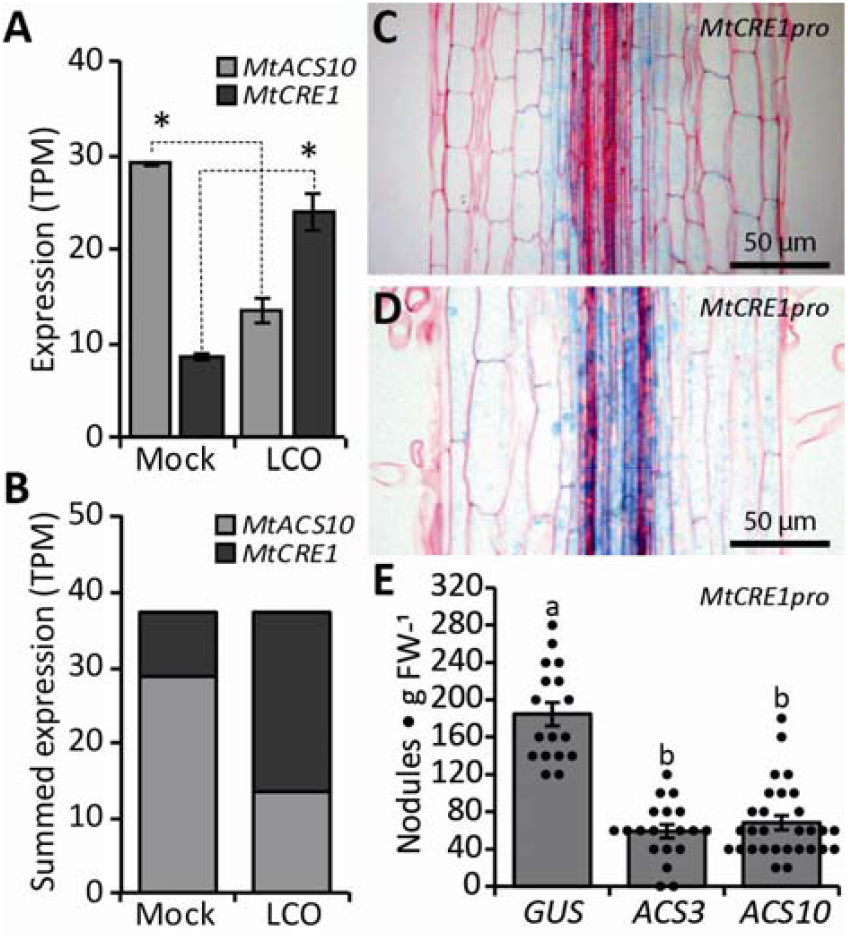
Counteracting *MtACS10* reduction and its effect on nodulation. (A-B) Antagonistic expression responses of *MtACS10* and *MtCRE1* under mock and lipo-chitooligosaccharide (LCO) treatment, shown as (A) individual and (B) summed expression bar graphs. Bars represent mean ± SE; n=3 (independent biological replicates each consisting of the susceptible zones of ∼16 pooled plate grown plants), an asterisk (*) indicate significant differences (Student t-test, *P* < 0.05). (C-D) *MtCRE1pro::GUS* reporter activity at 24 hours post-inoculation with (C) mock or (D) *Sm201*). (E) Effect of local ectopic induction of *ACS* gene expression on nodulation. Nodule numbers in control (*MtCRE1pro::GUS*) and two independent ectopic induced *ACS* expression lines (*MtCRE1pro::ACS3* and *MtCRE1pro::ACS10*) (n>15). Transgenic and non-transgenic roots (for untransformed roots see Supplemental Fig. S8) were distinguished by *DsRed* expression. Each dot represents an individual transgenic root (n>17); bars represent mean ± SE; different letters indicate statistical significance (one-way ANOVA followed by Tukey’s HSD test, *P* < 0.05). FW, fresh weight, ep, epidermis;c1-c5, cortical cell layers; en, endodermis; p, pericycle in C, D.

To test the functional relevance of the *MtACS10* repression, we generated composite plants bearing transgenic roots that ectopically express either *GUS* or *MtACS10* under control of the *MtCRE1* promoter. To rule out a gene-specific effect of *MtACS10*, we also created a construct in which *MtACS3* is driven by the *MtCRE1* promoter. The *MtCRE1pro::GUS* construct served both as a negative control and to validate that the promoter is active in the appropriate root cell layers (i.e. the pericycle, endodermis, and inner cortex, Fig. 7C) and is induced following LCO perception (Fig. 7D). As expected, expression of *MtCRE1pro::GUS* had no detectable effect on nodulation (Fig. 7E). In contrast, ectopic expression of either *MtACS3* or *MtACS10* under the *MtCRE1* promoter led to a 50– 60% reduction in nodulation (Fig. 7E). This effect was absent in the non-transgenic roots harvested from the same composite plants (Supplemental Figure S12). These findings demonstrate that temporal ectopic expression of either *ACS* gene in root interior represses nodulation.

### Ethylene helps determine the size of the Medicago root susceptible zone in apical-basal direction

As nodulation in Medicago seems to rely on the suppression of a local ethylene potential by a transcriptional repression of *MtACS10*, we next asked whether ethylene might also define the spatial limits of the root susceptible zone. To test this, we flooded the root of Medicago wild-type A17 and *sickle* root with *Sm2011* LCOs for 3 hrs and sampled the lower 2.5 cm of the root in five 5 mm segments. These segments were named from the root tip upwards as Z1 to Z5 (Fig. 8A). From these segments, we isolated RNA for cDNA synthesis and subsequent qRT-PCR against one of the key genes in nodulation, *NODULE INCEPTION (NIN)*. This revealed that in wild type, *NIN* induction is mostly restricted to Z2, the susceptible zone (Fig. 8B). We observed *NIN* induction in Z1 and Z3; however, these increases were marginal compared to the approximately 90-fold induction detected in Z2. This limited activation in the flanking zones may reflect natural variation in the position or extent of the susceptible zone, which can occasionally extend beyond the 5 mm segment defined as Z2. Similarly, the induction observed in Z4 suggests that the susceptible zone may sometimes expand further along the root or may not be strictly confined to a fixed location. In sickle, *NIN* was induced in all segments tested, although to a lesser extent in Z1 (Fig. 8B). This suggests that the response to LCO application is broader in this mutant. Next, we tested if nodulation could occur outside the susceptible zone. For this, we spot-inoculated wild-type A17 and *sickle* at different positions on the root (i.e. either Z1, Z2, Z3, Z4 or Z5) and counted the number of nodules at 7 DPI. In wild type, nodules were consistently formed on Z2, the susceptible zone, and Z1 (Fig. 8C), the latter probably reflecting the transition of Z1 into Z2 during the course of the experiment. This is not the case for the other zones. Apart from Z3, that only formed nodules occasionally, Z4 and Z5 never formed any nodules. When we performed this experiment in *sickle*, the results were different. In this mutant, nodules were formed on all zones, although marginally less efficiently in Z5 (Fig. 8C).

**Figure 8.**
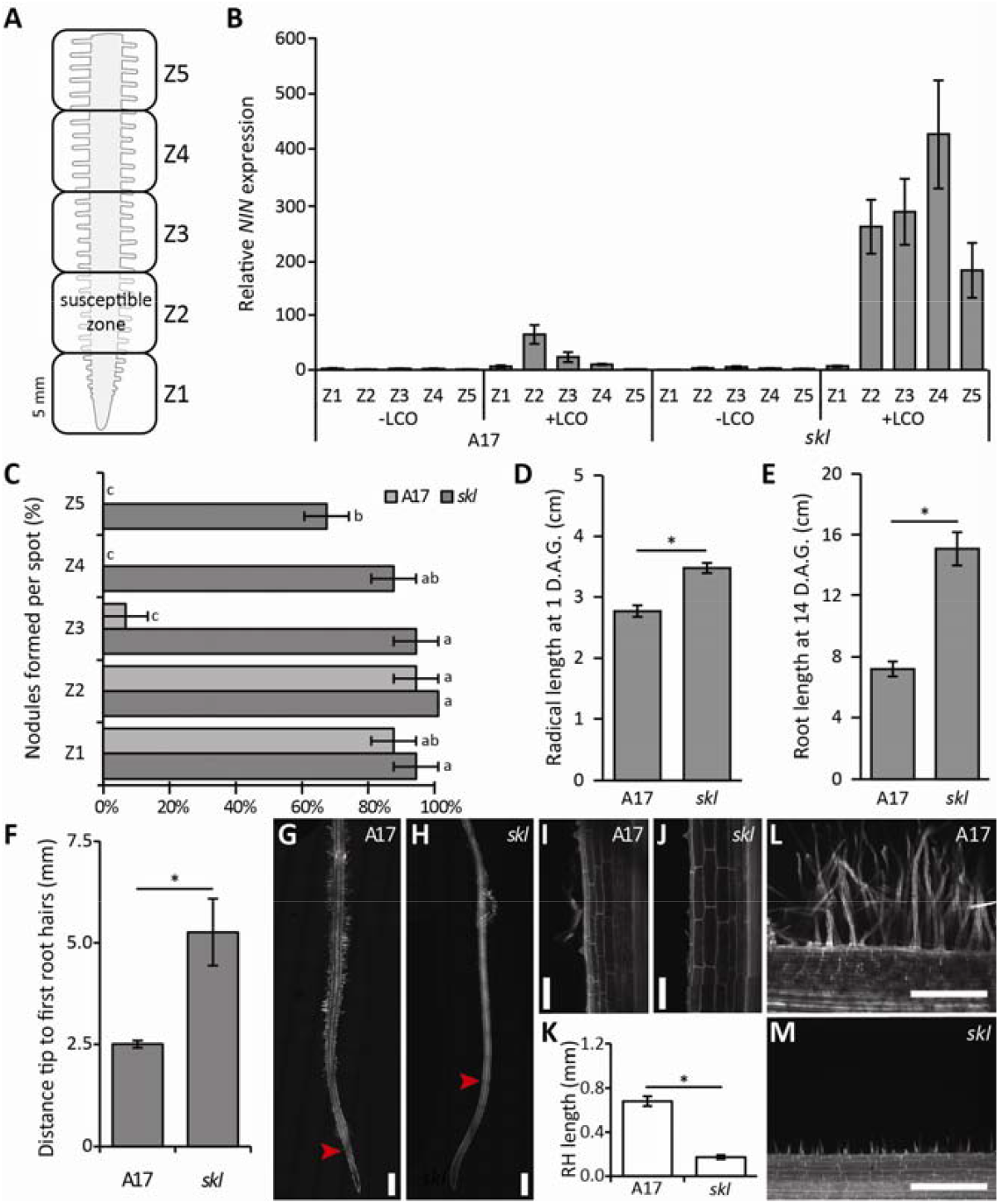
The role of ethylene in defining the root susceptible zone in Medicago. (A) Schematic representation of the Medicago root separation in 5 mm zones numbered bottom up, Zone (Z)1-5. (B) Relative *NIN* expression (n=3, independent biological replicates each consisting of the susceptible zones of ∼16 pooled plate grown plants) in Z1 to Z5 in response to Sm2011 lipo-chitooligosaccharide (LCO) in wild-type A17 and the *sickle(skl)/ethylene insensitive2* mutant of Medicago. (C) Percentage of nodule formed per spotted location at different distances from the root tip in wild-type A17 and *skl*. Bars represent mean ± SE; n>10, independent biological replicates each consisting of a spot-inoculation of the indicated root zone on plate grown plants; different letters indicate statistical significance (one-way ANOVA followed by Tukey’s HSD test, *P* < 0.05). Root length of wild-type A17 and *skl* at (D) 1 and (E) 14 days after germination (D.A.G.). Bars represent mean ± SE; n>15, independent biological replicates of a plate grown root; an asterisk (*) indicate significant differences (Student t-test, *P* < 0.05). (F) The average distance from the Medicago root tip where the first root hairs are formed in wild-type A17 and *skl*. Bars represent mean ± SE; n>5; an asterisk (*) indicate significant differences (Student t-test, *P* < 0.05). (G-H) Representative confocal tile image of the Medicago (G) wild-type A17 and (H) *skl* root tips. Arrowhead points at first root hair. (I-J) Zoom in of first root hairs in (I) wild-type A17 and (J) *skl*. (K) Average length of the root hairs (RH) in the susceptible zone measured at roughly 5 mm distance from the root tip in wild-type A17 and 10 mm in *skl*. Bars represent mean ± SE; n>6, multiple root hairs from minimally 6 plants measured per repicate; an asterisk (*) indicate significant differences (Student t-test, *P* < 0.05). (L-M) Representative confocal image of root hairs in the root susceptible zone of (L) wild-type A17 and (M) *skl*. Scale bars 1 mm in G, 100 µm in I and J, and 500 µm in L and M; n, independent biological replicates.

A key feature of the *sickle* mutant root is its length (Penmetsa and Cook 1997; Penmetsa et al. 2003). By 1 day after germination (DAG), the radical of the *sickle* mutant was approximately 25% longer than those of wild-type A17 (Fig. 8D). This difference in primary root length persisted and increased over time, reaching roughly a two-fold difference by 14 DAG (Fig. 8E). Additionally, we observed that the distance from the epidermal cells producing the first root hairs to the root tip was increased by roughly two-fold compared to wild-type A17 (∼2.5 vs 5.0 mm, Fig. 8F-J). Interestingly, *sickle* root hairs remained small and few over the entire length of the root susceptible zone, whereas in wild-type A17, root hairs became substantially longer within a short distance from their initiation (Fig. 8K-M). This phenotype was reminiscent of that observed in *Mtacs3-1* (Supplemental Figure S13A). Similar to sickle, root hairs in this mutant emerged further from the root tip and remained smaller than in wild-type R108, a characteristic not observed in the *Mtacs10-1* mutant (Supplemental Figure S13B-J).

We next examined the expression profiles of MtACS3 and MtACS10 along the developmental root axis. MtACS3 transcripts were predominantly detected in the root tip, with comparatively low expression in more proximal regions, whereas MtACS10 displayed an inverse pattern. Upon LCO treatment, induction of MtACS3 was largely confined to the root susceptible zone, while downregulation of MtACS10 extended beyond this region, including a significant but less pronounced reduction in the adjacent proximal zone (Supplemental Figure S13A,B).

To assess whether these spatial expression patterns correlate with ACC biosynthetic output, we quantified ACC levels in dissected root segments corresponding to the root tip and susceptible zone, both prior to and during LCO treatment. ACC levels were highest in the root tip and lower, with relatively uniform levels, in the remaining root regions. LCO treatment did not significantly alter ACC content in the susceptible zone (Supplemental Figure S13C). Collectively, these data establish a spatial relationship between root developmental dynamics, ACS gene expression, and ACC distribution along the root axis.

## DISCUSSION

This study provides insights into the spatial and functional dynamics of ethylene biosynthesis and signaling during the early stages of legume-rhizobium symbiosis in *Medicago truncatula*. By employing promoter–reporter constructs, mutant analyses, and transgenic approaches, we uncover a coordinated regulatory mechanism by which ethylene biosynthesis shapes the spatial domain and developmental outcome of nodule formation, with particular emphasis on the differential regulation and functional contributions of *MtACS3* and *MtACS10* (Fig. 9A-D). Our findings expand our understanding of how local ethylene biosynthesis and response contribute to the formation of nodules and infection threads, and their radial positioning (Fig. 9E)

**Figure 9.**
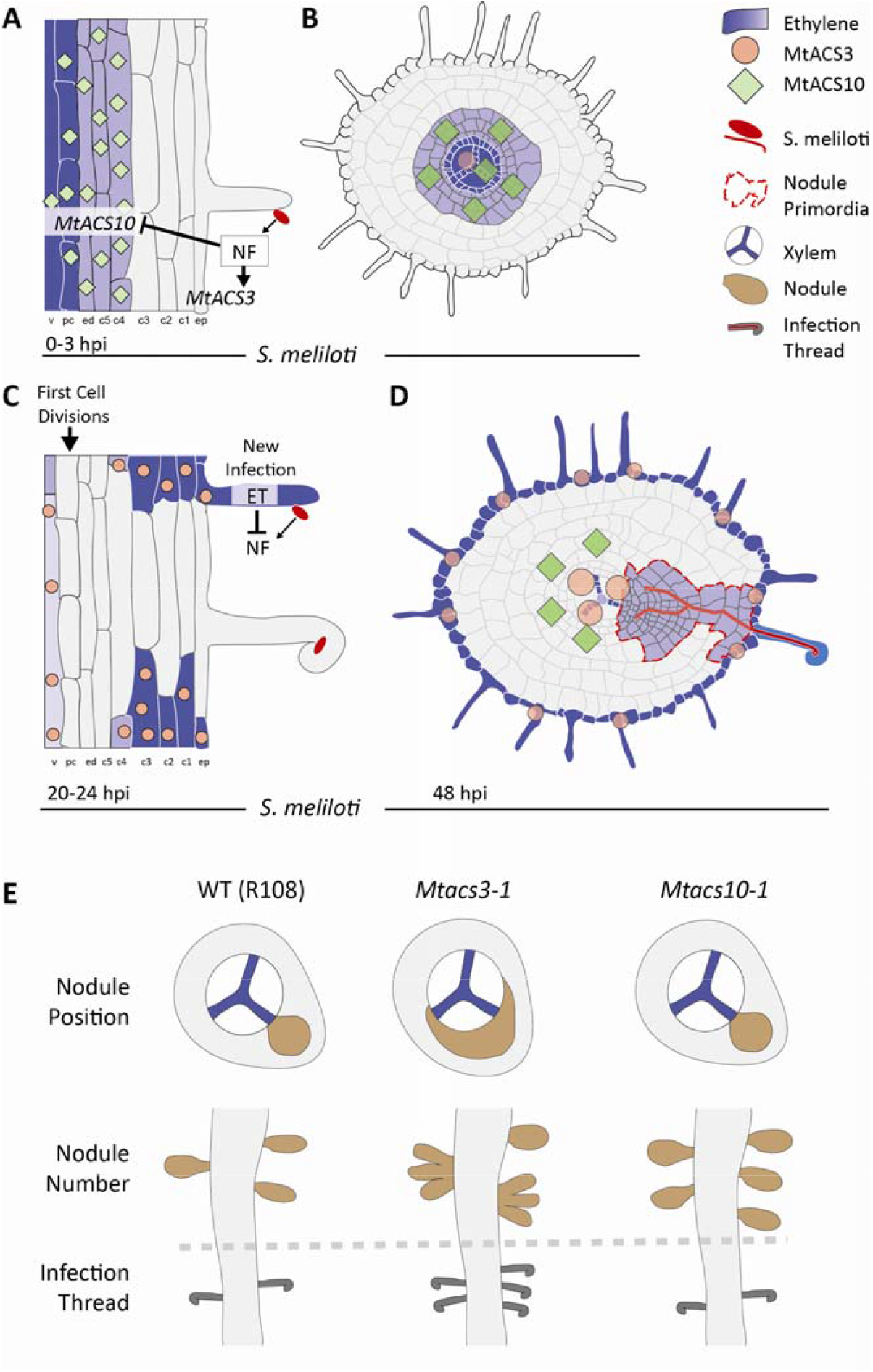
Proposed model of ethylene redistribution during nodule initiation in Medicago truncatula. (A, B) In the uninduced susceptible zone, ethylene is produced internally via MtACS10. Upon *Sinorhizobium meliloti* inoculation or Nod factor (NF) perception, *MtACS10* is suppressed while *MtACS3* is activated in the outer root layers, including the epidermis. (C, D) By 20–24 hpi, ethylene biosynthesis is shifted to the outer root tissues via *MtACS3*, possibly excluding the infected root hair, which inhibits additional root hair infection. At 48 hours post inoculation (hpi), the *EBSnew* ethylene reporter marks developing nodule primordia, with *MtACS3* proposed as a candidate contributing to this ethylene pulse. (E) Phenotypes of *Mtacs3-1* and *Mtacs10-1*: wild type (WT) and *Mtacs10-1* nodules remain xylem pole positioned, whereas this positional specificity is lost in *Mtacs3-1. Mtacs10* mutants form more nodules without changing the number of Infection Threads, while *Mtacs3* shows more infection threads, fused nodules, but a similar total number of nodulation events.

We first analyzed the spatiotemporal activity of the synthetic ethylene-responsive promoter *EBSn* during early rhizobial signaling. The initial localization of promoter activity in the inner tissues of the root susceptible zone under mock conditions likely reflects baseline ethylene biosynthesis, possibly through *MtACS10*. Upon rhizobial inoculation, however, *EBSn* activity shifts to the outer root layers, and eventually to the epidermis, suggesting either a redistribution of ethylene or, given our MtACS results, more likely a spatially restricted repression and induction of local ethylene biosynthesis. Given that ethylene is a gaseous molecule, one might expect diffusion to obscure such sharp expression boundaries. It is often assumed that ethylene can diffuse freely between cells and tissues (Park et al. 2017). Nevertheless, our observations show that localization of biosynthesis is crucial for nodulation. Ethylene biosynthesis occurs inside the cell and is perceived by receptors that are located on the endoplasmic reticulum (Chen et al. 2002; Wang et al. 2002; Ma et al. 2006; Ju and Chang 2015). This suggests that for ethylene to be functional, it has to be dissolved in the luminal fluid of the endoplasmic reticulum. The diffusion rate of dissolved ethylene in water (∼1–2 × 10^−5^ cm^2^ s^−1^) is three to four orders of magnitude lower than that of ethylene in air (∼1.4 × 10^−1^ cm^2^ s^−1^) (Huq and Wood 1968; Pritchard and Currie 1982). As such, it is likely that spatial separation of ethylene biosynthesis in different tissues can create concentration gradients, similar as is observed for other hormones (Friml et al. 2003; Grieneisen et al. 2007; Antoniadi et al. 2015; Rizza and Jones 2019). The localized changes in *MtACS* and *EBSn* promoter activity strongly argue for cell-type-specific biosynthesis, regulated transcriptionally. Shifting ethylene biosynthesis from inner to outer root tissues could also explain the increased ethylene emissions after rhizobia application that was previously reported (Ligero et al. 1986, 1987; Reid et al. 2018), even without changes in net ACC content, as ethylene is likely to leak more freely into the rhizosphere from the root epidermis.

Through RNA sequencing, qRT-PCR, and promoter–reporter analyses, we identified *MtACS3* and *MtACS10* as the predominant *ACS* genes expressed along the root developmental axis under non-symbiotic conditions, although with distinct spatial expression domains. RNA *in situ* hybridization further supported the spatial restriction of *MtACS10* to inner cortical domains. While *MtACS3* transcripts were not robustly detected in the susceptible zone by *in situ* hybridization, reporter analyses revealed inducible promoter activity upon rhizobial inoculation, underscoring the dynamic regulation of this gene during early symbiotic signalling. Together, the distinct and spatially regulated expression patterns of *MtACS3* and *MtACS10* suggest a mechanism by which ethylene biosynthesis is locally reconfigured to balance infection control with nodule initiation.

The functional relevance of these transcriptional shifts is highlighted by RNAi and mutant analyses. Knockdown or loss-of-function of *MtACS10* results in a significant increase in nodule number, suggesting that this gene acts as a local suppressor of nodulation. Its downregulation after LCO treatment appears to be a critical step in rendering specific pericycle and/or cortical cells competent for division and nodule primordium formation. This interpretation is supported by the finding that ectopic expression of either *MtACS3* or *MtACS10* under the *MtCRE1* promoter represses nodulation. This supports the hypothesis that downregulation of *MtACS10*, and likely ethylene biosynthesis, in specific cell layers is a prerequisite for successful nodule initiation in Medicago. By preventing this repression, the developmental program is hindered, reinforcing the functional importance of spatial gene regulation. Conversely, MtACS3 likely affects nodule number indirectly by influencing infection thread frequency and, when mutated, promotes nodule clustering. As we expected this response to be regulated from the inner root tissues, the *MtACS3* expression domain indicates the observed nodule clusters might be the result of secondary infections. This would suggest that MtACS3 fine-tunes the infection process and possibly spatial coordination among neighboring infection events, potentially by modulating ethylene levels in the epidermis. The induction of *MtACS3* expression in the root hairs observed by Breakspear at al. (2014) supports this latter statement. We hypothesize that an epidermal increase in ethylene levels in response to LCO perception is part of a negative feedback loop that restricts further signalling after a successful symbiotic interaction has been initiated. This hypothesis is consistent with the hyper-infection phenotype observed in the sickle mutant (Penmetsa and Cook, 1997). The induction of additional *ACS* family members (e.g. *MtACS1* and *MtACS8*) following rhizobial application further suggests that *MtACS3* likely functions as part of a broader ethylene biosynthetic network rather than acting alone at this stage.

Additionally, our analyses indicate that epidermal infection thread formation in *Mtacs10-1* is comparable to that observed in R108, suggesting that MtACS10 does not substantially influence infection initiation at the epidermal level. However, nodule formation is a multistep process that extends beyond early epidermal infection and likely involves additional regulatory checkpoints in inner cortical and pericycle tissues that determine whether an initiated infection progresses into a mature nodule. Because infection thread quantification captures a defined developmental stage, it does not necessarily predict final nodulation outcome. It is therefore conceivable that, in wild-type plants, only a subset of initiated infections successfully transition into nodules, whereas in *Mtacs10-1* a greater proportion of these events proceed through later developmental checkpoints. Suggesting a role for *MtACS10*, and likely ethylene, at this stage as well.

Importantly, the radial positioning of nodule primordia, a process previously shown to be ethylene-sensitive in pea and Medicago (Heidstra et al. 1997; Penmetsa et al. 2003, 2008), is altered in the *Mtacs3-1* mutant. Wild-type Medicago roots preferentially form nodules opposite xylem poles, a pattern disrupted in *Mtacs3-1* where nodule initiation is more randomly distributed. These results suggest that local ethylene biosynthesis via *MtACS3* contributes to defining or reinforcing polarity cues for nodule positioning. We cannot exclude that the observed vasculature promoter activity upon rhizobial inoculation is sufficient to create a local restriction on nodulation near the phloem poles. Nevertheless, even if a vasculature-associated expression domain of *MtACS3* would be sufficient to direct nodule initiation toward the xylem poles, the fact that *MtACS3* promoter activity is only observed after rhizobial application raises questions on what determines the position of this local ethylene biosynthesis. Our results suggest that ethylene produced via MtACS3 may not establish positional information itself but instead acts downstream of an unknown cue that determines the radial bias. However, in the *Mtacs3* mutant, some nodules seem to spread wider after initiation, whereas at other nodule primordia, infection threads that originate in the epidermis opposite the phloem poles are clearly observed. This suggests that epidermal ethylene biosynthesis could indirectly co-regulate nodule numbers and taxis and indicates that secondary infections in the *Mtacs3-1* mutant lead, at least in part, to the observed disruption in radial nodule positioning. Interestingly, although *MtACS10* is expressed in the pericycle and inner cortex, *Mtacs10-1* mutants retain wild-type spatial bias, reinforcing the notion that MtACS10 primarily functions in restricting nodule number rather than spatial placement.

The ethylene-insensitive *sickle* mutant demonstrates that ethylene not only regulates local infection and nodule formation but also co-defines the spatial extent of the root susceptible zone. In wild-type roots, the LCO-induced expression of *MtNIN* is largely confined to a narrow region roughly 5 mm distal from the root tip, while in *sickle*, this response is, with the exception of the root tip, broadened to the entire 20 mm root segment analyzed. This expansion is consistent with the broader nodulation capacity of *sickle* roots, which form nodules outside the conventional susceptible zone. Additionally, the altered root growth dynamics in *sickle*, including extended elongation zone and altered root hair development, underscore the broader developmental roles of ethylene that indirectly influence symbiotic competence.

Ethylene serves as a central developmental “timer” in plant tissues, promoting their maturation and eventual senescence (Khan et al. 2014). Additionally in Arabidopsis, ethylene also enhances root hair elongation by modulating key regulatory pathways (Pitts et al. 1998). In line with this, our findings suggest that MtACS3 may act as a key regulator of root hair maturation and infection competence in Medicago. RNA *in situ* hybridisation shows that *MtACS3* is expressed in the epidermis at the start of the differentiation zone, precisely where root hairs are emerging. This is consistent with the observation that in the *Mtacs3-1* mutant, root hairs remain small and poorly developed, a phenotype strongly reminiscent of the ethylene-insensitive *sickle/Mtein2* mutant. Although small and poorly developed, the root hairs of *Mtacs3-1* still support rhizobial infection, indicating that ethylene is not strictly required to establish rhizobia susceptibility. Instead, we suggest that ethylene regulates the duration of the susceptibility window. When root hairs emerge at the start of the differentiation zone, ethylene may initiate a physiological timer that defines both the temporal and spatial extent of the susceptible zone. In this framework, wild-type root hairs in the susceptible zone remain competent for rhizobial entry, whereas those positioned beyond it have matured past susceptibility. A rhizobia-induced burst of ACS3-derived ethylene in the epidermis upon inoculation could then accelerate maturation of neighbouring, non-infected hairs, effectively pre-closing the infection window. In this model, a strong upregulation of *MtACS3* would function as a switch—allowing infection of a limited number of hairs while shielding the root from undesired supernumerary infections. In the absence of *MtACS3* activity, or if ethylene signalling is compromised, as in *Mtacs3-1* or *sickle/Mtein2*, respectively, both maturation controls (i.e., physiological timer and switch) are lost, leading to prolonged susceptibility, excess infection threads, and clustered nodules. Thus, ACS3-mediated ethylene production may function as a que that synchronizes the transition from susceptibility to resistance at the root surface.

Taken together, these findings establish a new framework in which ethylene signaling, mediated by distinct *ACS* genes, functions in a spatially and temporally resolved manner to balance infection thread formation, restrict excessive nodulation, and co-defines the physical limits of the root susceptible zone. The selective repression of *MtACS10* emerges as a key step in nodule initiation in Medicago, while *MtACS3* modulates infection-related parameters. This dual regulation of ethylene biosynthesis, both activation and repression depending on tissue context and developmental timing, illustrates once more the sophistication of hormonal control in root nodule symbiosis. Moreover, this dynamic regulation of ethylene biosynthesis provides insight into how plants use spatial and functional partitioning of gene activity to integrate multiple signaling layers to fine-tune developmental responses like nodulation with precision. For legume-rhizobia interactions, this spatial specificity allows the plant to limit the cost of nodulation by tightly controlling where and how nodules form. Future work should test whether similar regulatory mechanisms operate in other legumes, particularly crops, and how autoregulation of nodulation or environmental factors such as nitrate and abiotic stress modulate this ethylene-dependent spatial control.

To date, engineering nodulation in non-legumes has largely emphasized activating positive regulators, while inhibitory pathways remain underexplored (Untergasser et al. 2012; Mus et al. 2016; Huisman and Geurts 2020). Our findings suggest that ethylene homeostasis in Medicago restricts pericycle and cortical divisions, mirroring its conserved role in lateral root initiation in *Arabidopsis* (Nodzon et al. 2004; Prasad et al. 2010). Given the proposed shared origin of nodules and lateral roots (Nutman 1948; Hirsch et al. 1997; Mathesius et al. 2000; Franssen et al. 2015; Schiessl et al. 2019; Soyano et al. 2019), ethylene or similar repressors may likewise limit nodule initiation in other species. Accounting for such inhibitory mechanisms will be essential for engineering nodulation in non-legume crops.

## MATERIALS AND METHODS

### Plant material, growth conditions and treatments

Medicago seeds (wild-type Jemalong A17 and R108, *Mtacs3-1* (NF0616), *Mtacs3-2* (NF1952),*Mtacs10-1* (NF15254) and Mtacs10-2 (NF2329)) were treated with concentrated H_2_SO_4_ for 7 minutes, rinsed five times with MilliQ water, and sterilized for 10 minutes using normal household bleach. Seeds were again washed with sterile MilliQ water five times and placed on round Petri dishes containing Fåhraeus medium (0.25 mM NO_3_ ^-^, 1% Daishin agar, 0.01% (w/v) carbendazim) at 4°C in darkness for stratification. After 48 hours, seeds were transferred to room temperature to germinate for an additional 24 hours. Germinated seedlings were transferred to the appropriate growth system. For all experiments, plants were grown in an environmentally controlled growth chamber at 20°C/18°C with a 16h-light/8h-dark cycle and 70% relative humidity. A modified Fåhraeus medium (0.12 g/L MgSO_4_•7H_2_O, 0.10 g/L KH_2_PO_4_, 0.15 g/L Na_2_HPO_4_•2H_2_O, 1 ml/L 15 mM Fe-Citrate, 2.50 ml/L Spore-elements β- (CuSO_4_•5H_2_O 0.0354g/L, MnSO_4_•H_2_O 0.462g/L, ZnSO_4_•7H_2_O 0.974g/L, H_3_BO_3_ 1.269 g/L, Na_2_MoO_4_•2H_2_O 0. 398 g/L), pH 6.7) (Fähraeus 1957) with 0.125 mM Ca(NO_3_)_2_ was used in all experiments. For square (12cmx12cm) plate grown plants, 1% Daishin agar was added. Eight seedlings were grown per plate, and four plates per were used per replicate. All plates were partially covered with tin foil to avoid light grown roots. From the growing roots, the root susceptible zones (± 0.5 cm) of ca. 20-30 roots were collected and snap frozen in liquid nitrogen. All samples were stored at −80 °C until further use.

### LCO or Rhizobia application

*Sinorhizobium meliloti strain 2011* (Sm2011) LCOs were purified and applied as previously described (Spaink et al. 1991; van Zeijl et al. 2015b) with minor modifications. Sm2011 LCO stocks were stored in 100% DMSO and diluted 100-fold (∼10-9 M) in Fåhraeus (0.125 mM Ca(NO_3_)_2_) medium prior to application. The same Fåhraeus medium with 1% DMSO was used as mock treatment. LCO or mock treatments were pipetted on the root susceptible zone. Roots were exposed for 3 hours and subsequently the root susceptible zone (0.5 cm root segments) were cut just 5 mm from the root tip and snap-frozen in liquid nitrogen (n=3) for gene expression analysis. Sm2011 was grown on liquid YEM medium, spined down and the pallet resuspended in Fåhraeus medium to an OD_600_ of 0.02 or 0.1 for direct application on either plates or inoculation in plants grown on perlite, respectively.

### Phylogenetic analyses

ACS, ACO, and ACD protein sequences were retrieved from various plant genome databases, including that of Medicago (see supplemental table S1), using the BLASTP algorithm (Altschul et al. 1990). Homologous sequences were aligned using MAFFT version 7.450 with automatic selection of the appropriate algorithm and using the BLOSUM62 scoring matrix (Henikoff and Henikoff 1992; Katoh et al. 2002). Phylogenetic trees were estimated based on maximum likelihood using the IQTree webserver version 1.6.11 (Trifinopoulos et al. 2016; Minh et al. 2020) with automatic selection of the best-fitting model of protein evolution based on BIC using ModelFinder (ACS and ACO: JTT+F+I+G4; ACD: LG+G4 (Kalyaanamoorthy et al. 2017)) and 1000 ultrafast bootstrap replicates to assess clade support (Hoang et al. 2017).

### RNA isolation, cDNA synthesis and Quantitative RT-PCR

RNA was isolated from snap-frozen roots samples using the plant RNA kit (E.Z.N.A, Omega Biotek, Norcross, USA) according to the manufacturer’s protocol. This RNA was either send for sequencing (Beijing Genomics Institute, Hong Kong, China) or 1 μg total RNA was used to synthesize cDNA using the i-script cDNA synthesis kit (Bio-Rad, Hercules, USA) as described in the manufacturer’s protocol. Real time qRT-PCR was set up in 10 µl reactions with 2× iQ SYBR Green Super-mix (Bio-Rad, Hercules, USA). Experiments have been conducted on a CFX Connect optical cycler, according to the manufacturers protocol (Bio-Rad, Hercules, USA). All primers including the genes used for normalization (*MtUBQ10* and *MtPTB*) are given in Supplemental table S2. Data analysis was performed using CFX Manager 3.0 software (Bio-Rad, Hercules, USA). Cq values of 32 and higher were excluded from the analysis, though still checked for transcriptional induction. Statistical significance was determined based on student’s t-test (p<0.01).

### RNA Sequencing

RNA was sequenced at BGI Tech Solutions (Beijing Genomics Institute, Hong Kong, China) using the Illumina Truseq (Transcriptome) protocol utilizing an Illumina Hiseq 2000 instrument generating 150 bp paired-end reads. In total 24–33 million clean reads were generated for each sample. Sequencing data were analyzed by pseudoaligning RNA-seq reads against the *Medicago truncatula* genome annotation (Mt4.0v1) using kallisto v0.46.2 (Bray et al. 2016) with default settings. All reads are available on the European Nucleotide Archive (ENA) under accession number PRJEB38785.

### RNA in situ hybridisation

RNA in situ hybridisation was performed as previously described (Kulikova et al. 2018). RNA ISH probe set for *MtACS3* and *MtACS10* was designed and synthesized by request at Thermo Fisher Scientific (catalogue numbers VF1-6000770 and VF1-6000771 for *MtACS3* and *MtACS10*, respectively). As a negative control any probe set was omitted for hybridization. Probe IDs are given in Supplemental table S3.

### Vectors and constructs

For RNAi-mediated knockdown of *MtACS3* or *MtACS10*, two fragments (332 bp and 310 bp, respectively (Supplemental table S4)) were amplified from Medicago Jemalong A17 root cDNA, using specific primer pairs (Supplemental table S2), and cloned into pENTR-D-TOPO (Invitrogen, Carlsbad, USA). Both RNAi fragments were recombined into the DsRed-modified gateway vector pK7GWIWG2(II)-RR driven by the *CaMV35S* promoter (Limpens et al. 2004) to obtain the binary constructs pK7GWIWG2(II)-RR-*p35S-MtACS3*-RNAi and pK7GWIWG2(II)-RR-*p35S-MtACS10*-RNAi. For the empty vector control, the binary plasmid pK7GWIWG2(II)-RR-*p35S*-RNAi-control as previously described (van Zeijl et al. 2015a) was used. For ectopic expression and promoter GUS studies, the coding sequences, putative promoters (3.5 kb region upstream of start codon) and putative 3’UTRs (1 kb region downstream of stop codon) of *MtACS3* and *MtACS10* were synthesized at Thermo Fisher Scientific. These, as well as a GUS-encoding sequence (pICH75111), MtCRE1 promoter (Gonzalez-Rizzo et al. 2006) and *35S* terminator (pICH41414) were assembled together with the into vectors pICH47811 to create EC74831; MtCRE1p-MtACS3-t35S; EC74832; MtCRE1p-MtACS10-t35s; EC74833; MtCRE1p-GUS-t35s; EC74963; MtACS3p-GUS-ACS3-3`UTR; EC74964; MtACS10p-GUS-ACS10-3`UTR (for sequences see Supplemental table S5) using Golden Gate cloning (Engler et al. 2014). Binary transformation constructs were created by assembling the resulting clones together with clone pICH47732-*AtUBQ10::DsRED::tNOS* into vector pICSL4723 (Engler et al. 2014). All constructs are available from our laboratory upon request.

### Plant transformation and histology

*Agrobacterium rhizogenes*-mediated root transformation was used to transform Medicago (Jemalong A17) as previously described (Limpens et al. 2004). Transgenic roots were selected based on DsRED1 expression. Plants with transgenic roots were, depending on their use, transferred to either plates or perlite containing Fåhraeus medium (0.125 mM Ca_2_(NO_3_)_2_), and inoculated with the *S. meliloti strain 2011* (Sm2011, OD_600_ = 0.1) between 20-48 hours for most GUS work and two-three weeks in the case of nodulation assays after transfer.

### ACC extraction, detection and quantification by liquid chromatography-tandem mass spectrometry

ACC analysis from the Medicago root were performed as previously described (Bours et al. 2013) with adaptations as described by Gühl and colleagues (Gühl et al. 2021).

### Statistical Analysis

When appropriate, data were subjected to the Student’s t test (Microsoft Excel). All other data were subjected to one-way ANOVA. Individual differences were then identified using a post hoc Tukey test (P < 0.05). For these, all analyses were performed using SAS_9.20 (http://www.sas.com/).

## AUTHOR CONTRIBUTIONS

Conceptualization, SM, AvZ, WK; Methodology, RvV, OK, RH, WK; Investigation, SM, TS, KA, RH, AvS, RvV, OK, TW, JK, HF, EL, AvZ, WK; Formal Analysis, SM, TS, AvZ and WK; Contribution of a new analytic tool; JFM, AS, JA; Visualization, SM, WK; Writing – Original Draft, SM, TS, AvZ and WK; Writing – Review & Editing, SM, TS, AvS, RvV, OK, RH, JFM, AS, JA, HF, EL, AvZ, WK; Writing – Final version, SM, TS, AvZ and WK; Funding Acquisition, WK; Supervision, WK.

## ACKNOWLEDGMENTS

We would like to acknowledge Marijke Hartog and Sidney Linders for technical support of this project. Colleen Drapek for suggestions and critically reading of the manuscript.

