## Supplemental figures and tables for "Spatiotemporal dynamics of ethylene biosynthesis shape infection and nodule initiation in Medicago truncatula"

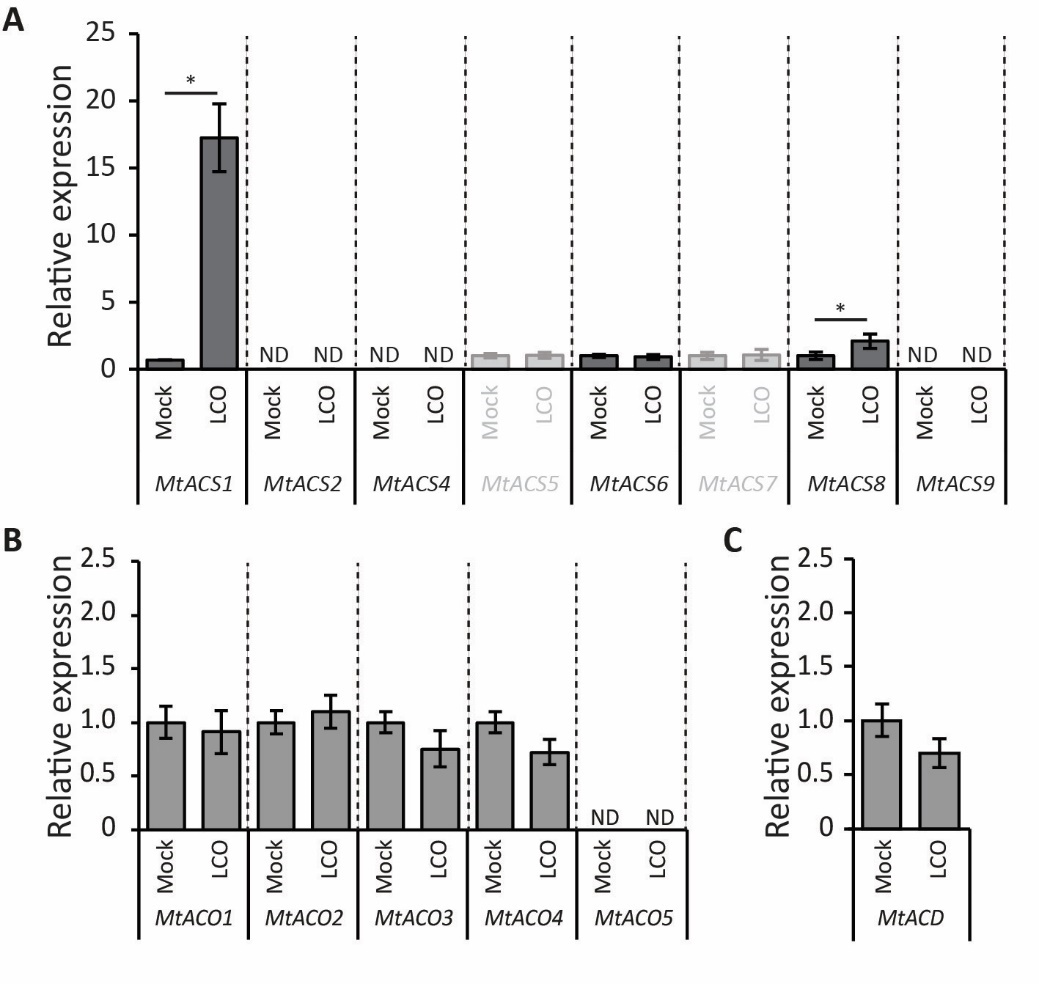

**Supplemental Figure S1.** Relative expression of ethylene biosynthesis and ACC metabolism gene following mock and lipo-chitooligosaccharide (LCO) application, measured at 3 hours post-treatment of (A) MtACS1, 2, 4, 5, 6, 7, 8, 9, (B) MtACO1, 2, 3, 4, 5, and (C) MtACD. Bars represent mean ± SE (n=3, independent biological replicates each consisting of the susceptible zones of ~16 pooled plate grown plants). Asterisks (*) indicate significant differences between treatments (Student t-test, P < 0.05).

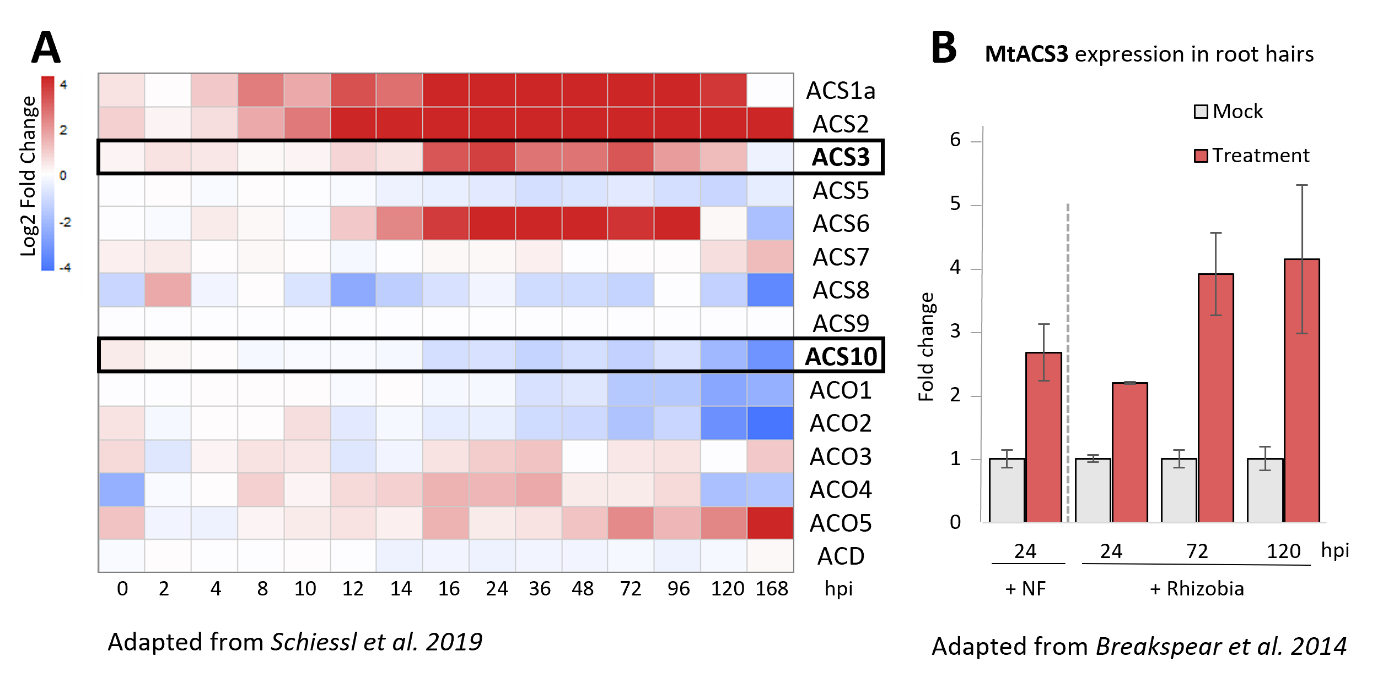

**Supplemental Figure S2.** Ethylene biosynthesis gene expression from published datasets. For reference, the gene names were unified according to Supplemental Table S1. (A) From Schiessl et al., (2019).(B) From Breakspear et al., (2014).

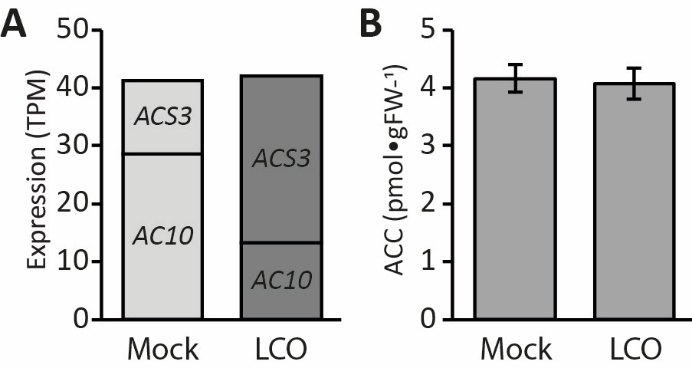

**Supplemental Figure S3:** The effect of 3 hours lipo-chitooligosaccharide (LCO) application on **(A)** cumulative *MtACS3* and *MtACS10* expression (TPM, transcripts per million), and **(B)** ACC levels. All bars represent means ± SE, (n=6); For B, no statistical significance differences (one-way ANOVA followed by Tukey’s HSD test, *P* > 0.05); n, 10-20 pooled root susceptible zones as independent biological replicates.

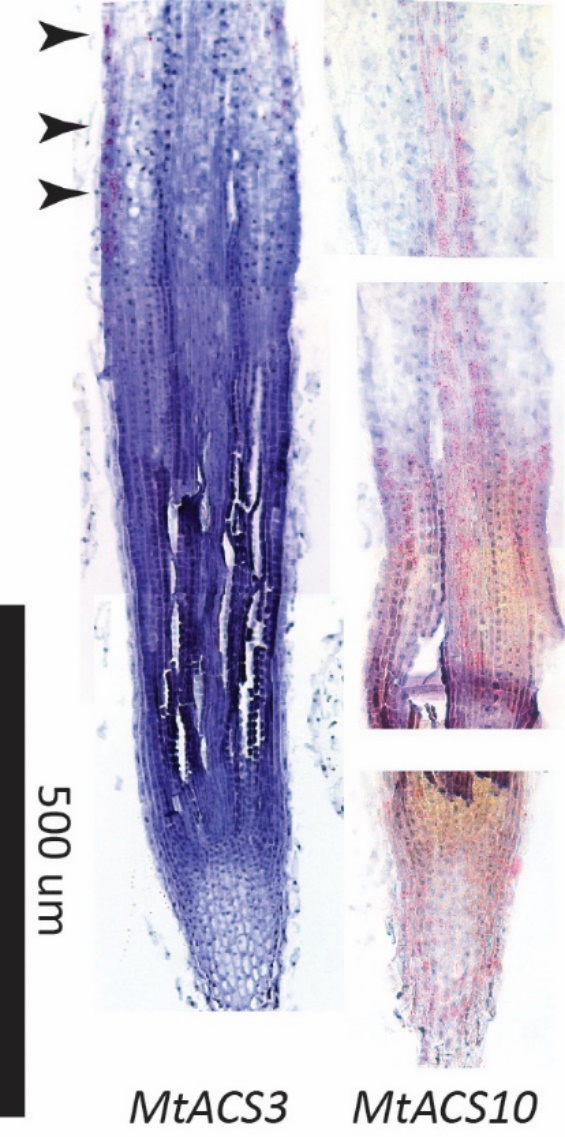

**Supplemental Figure S4.** Longitudinal sections of the untreated Medicago A17 wild-type root tip with *MtACS3* or *MtACS10* expression pattern visualized via RNA *in situ* hybridization. Hybridization signals appear as red dots (arrowheads highlight distinct expression domain of *MtACS3* in the epidermis and c1 layer of the start of the differentiation zone).

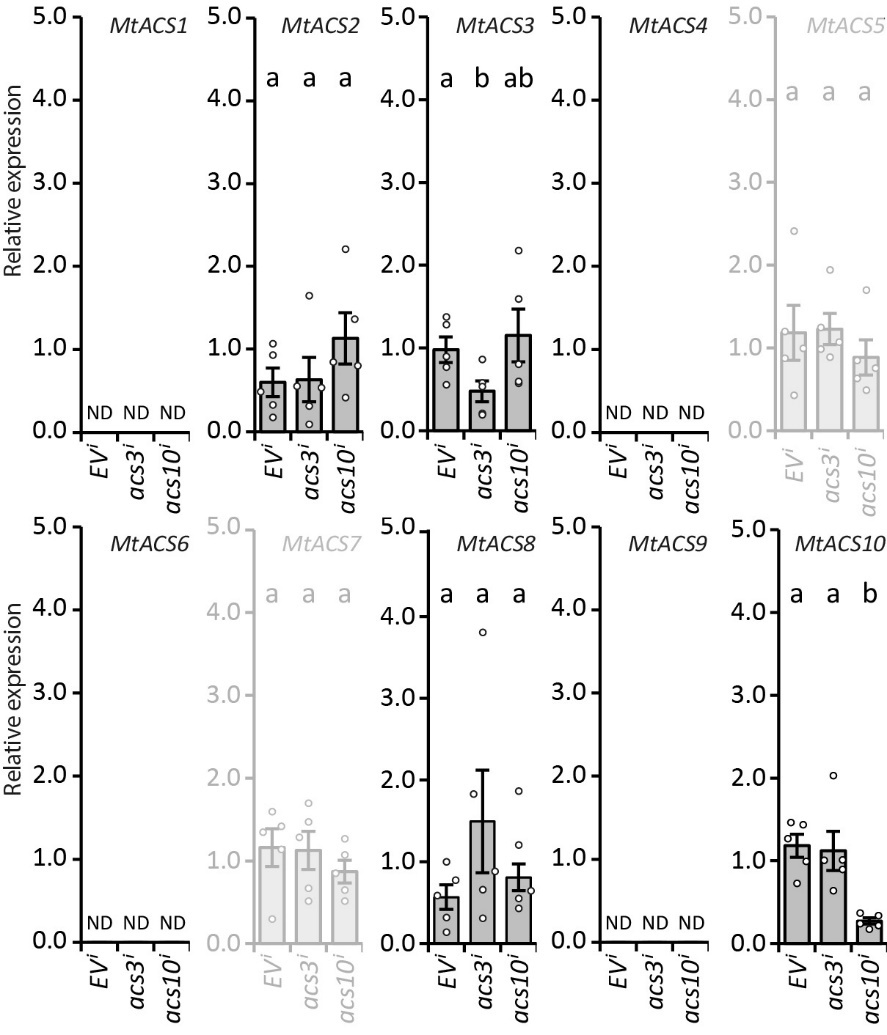

**Supplemental Figure S5:** Silencing and off-target effects of *ACS3^i^* and *ACS10^i^* on *MtACS1*, *MtACS2*, *MtACS3*, *MtACS4*, *MtACS5*, *MtACS6*, *MtACS7*, *MtACS8*, *MtACS9*, and *MtACS10* expression under mock conditions. Each dot represents an individual transgenic root; bars show mean ± SE Different letters indicate statistical significance between the Empty Vector (*EV^i^*) and *ACS3^i^* or *ACS10^i^* line (n=5, one-way ANOVA followed by Tukey’s HSD test, *P* < 0.05). Putative aminotransferases previously misidentified as ACC synthases in grey; ND, not detected; n, independent biological replicates.

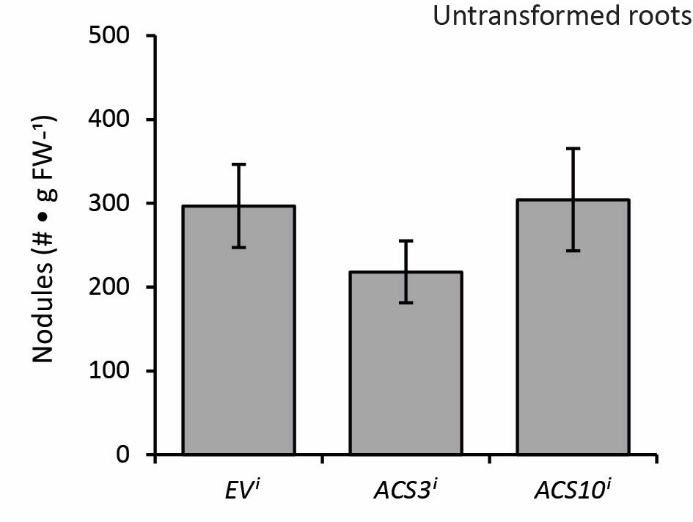
**Supplemental Figure S6:** Numbers of nodules formed on non-transgenic roots belonging to the composite plants of the Empty Vector (*EV^i^*) control and the two RNA interference (*ACS3^i^* and *ACS10^i^*) lines (n>10). Bars represent means ± SE. Non-transgenic roots were harvested based on absence of *DsRed* expression. Bars represent means ± SE. Non-transgenic roots were harvested based on absence of *DsRed* expression. No statistical significance differences (one-way ANOVA followed by Tukey’s HSD test, *P* > 0.05); n, non-transgenic roots from individual plants as independent biological replicates.

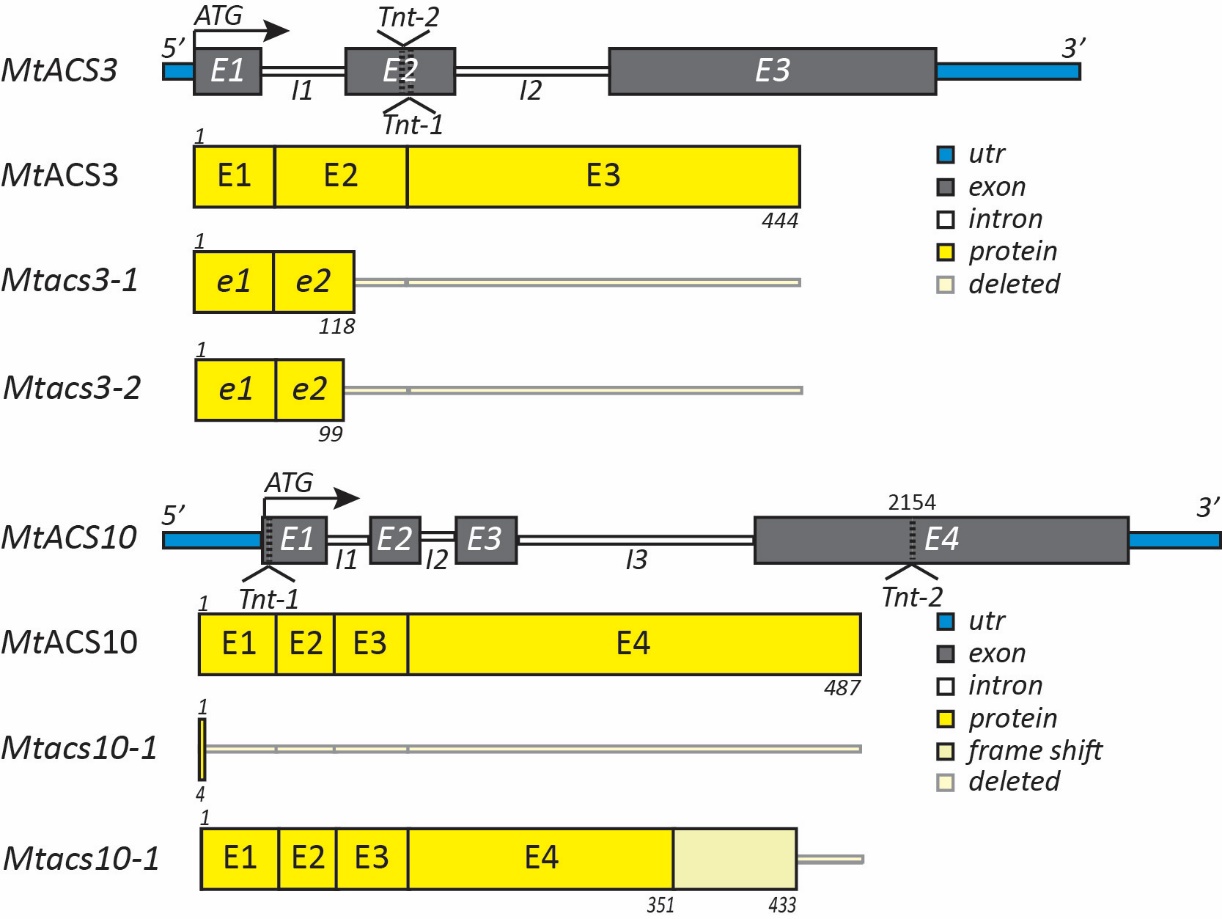

**Supplemental Figure S7:** *Tnt1* insertion mutations in *MtACS3* and *MtACS10*. Schematic representation of the gene structures of *MtACS3* (top) and *MtACS10* (bottom), showing untranslated regions (blue), exons (grey), introns (white), and predicted protein-coding regions (yellow). Positions of *Tnt1* retrotransposon insertions are indicated by triangles. The *Mtacs3-1* and *Mtacs3-2* allele carry an insertion in exon 2, resulting in a truncated predicted protein of 118 and 99 amino acids, respectively, compared to the full-length 444 aa MtACS3. The *Mtacs10-1* and *Mtacs10-2* alleles carry an insertion in either exon 1 and exon 4. The *Tnt1* insertion in *Mtacs10-1* leads to a severely truncated predicted protein of only 4 amino acids compared to the 487 aa full-length MtACS10, whereas *Mtacs10-2* leads to a frame shift at amino acid 351 followed by a premature stop codon at amino acid 433. Deleted regions are indicated in light grey.

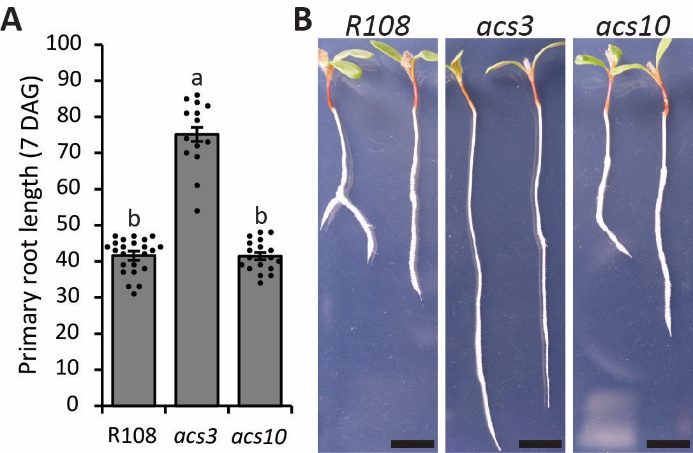
**Supplemental Figure S:** Loss of *MtACS3*, but not *MtACS10*, increases primary root length in Medicago. **(A)** Quantification of primary root length at 7 days after germination (DAG) in wild-type R108, *Mtacs3*, and *Mtacs10* mutant seedlings (n>14). Each dot represents an individual plate grown root; bars show mean ± SE Different letters indicate statistical significance (one-way ANOVA followed by Tukey’s HSD test, *P* < 0.001 for *Mtacs3* vs. R108 and *Mtacs3* vs. *Mtacs10*; n.s., not significant for *Mtacs10* vs. R108). **(B)** Representative seedlings of R108, *Mtacs3*, and *Mtacs10* at 7 DAG. Scale bars, 1 cm.

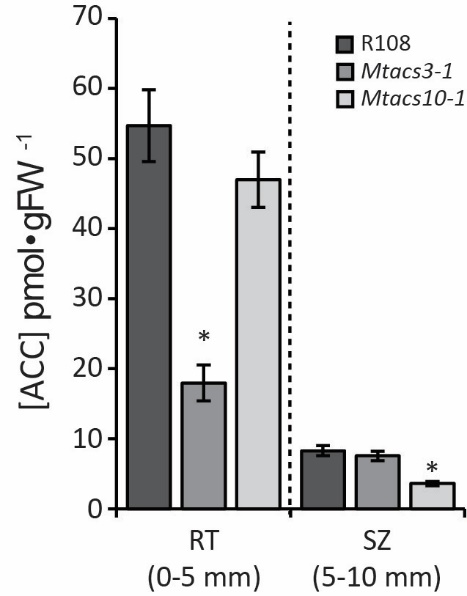
**Supplemental Figure S9:** Concentrations of ACC in R108 per gram FW in wild type, *Mtacs3-1* and *Mtacs10-1* measured at 0-5 mm and 5-10 mm from the root apex, root tip (RT) and susceptible zone (SZ) respectively. Bars represent mean ± SE; n=5, independent biological replicates each consisting of ~24 pooled plate grown root tips separated into two 5 mm segments; different letters indicate statistical significance (paired Student t-test, P < 0.05).

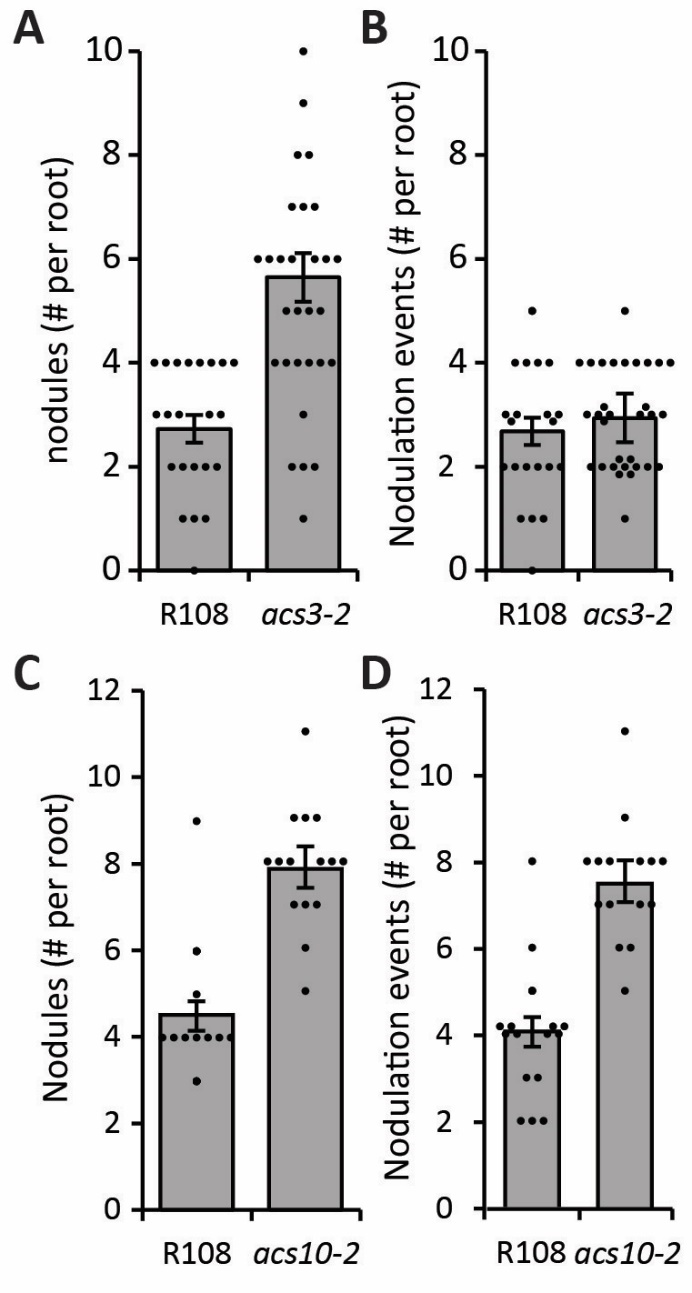

**Supplemental Figure S10:** Effect of *acs3-2* and *acs10-2* knockout mutations on nodulation in R108 plants grown on plates. (A) Average number of root nodules formed on R108 and *Mtacs3-2* mutants (n>22, independent roots of plate grown plants). (B) Number of nodule initiation sites per root (n>22, independent roots of plate grown plants, same plants as in A). (C) Average number of root nodules formed on R108 and *Mtacs10-1* (*acs10-2*) mutants (n>15, independent roots of plate grown plants). (D) Number of nodule initiation sites per root (n>15, independent roots of plate grown plants, same plants as in C). Each dot represents an individual root, bars represent mean ± SE; an asterisk (*) indicate significant differences (Student t-test, *P* < 0.05).

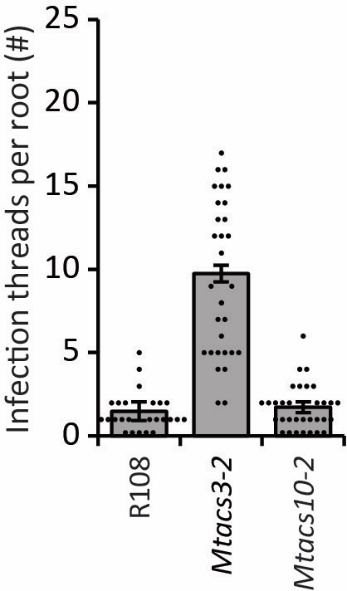
**Supplemental Figure S11:** Effect of *Mtacs3-2* and *Mtacs10-2* loss-of-function mutations on infection thread formation. Quantification of infection threads per spot-inoculation site following inoculation with GFP-labeled *Sm2011* (Sm2011-GFP) on wild-type R108, *Mtacs3-2*, and *Mtacs10-2*. Each dot represents an individual spot inoculated susceptible zone (n>24); bars represent mean ± SE; different letters indicate statistical significance (one-way ANOVA followed by Tukey’s HSD test, *P* < 0.05).

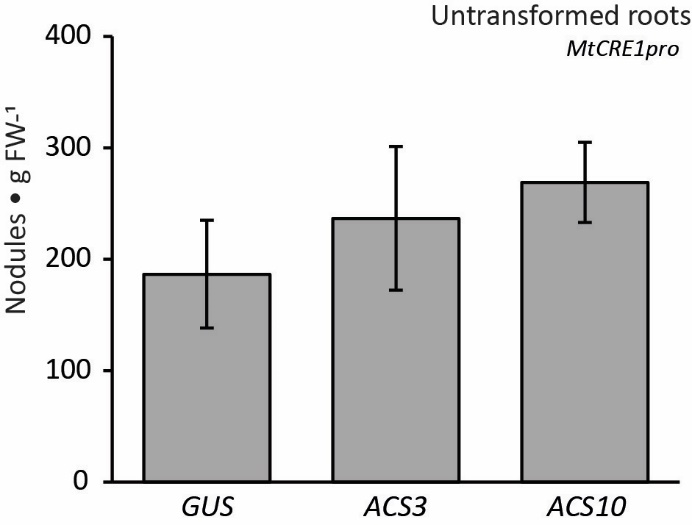
**Supplemental Figure S12:** Numbers of nodules formed on non-transgenic roots belonging to the composite plants of the control *GUS* (*MtCRE1pro::GUS*) and two independent ectopically induced *ACS* expression lines, *ACS3* (*MtCRE1pro::ACS3)* and *ACS10* (*MtCRE1pro::ACS10*) (n>15). Bars represent means ± SE. Non-transgenic roots were harvested based on absence of *DsRed* expression. No statistical significance differences (one-way ANOVA followed by Tukey’s HSD test, *P* > 0.05); n, non-transgenic roots from individual plants as independent biological replicates.

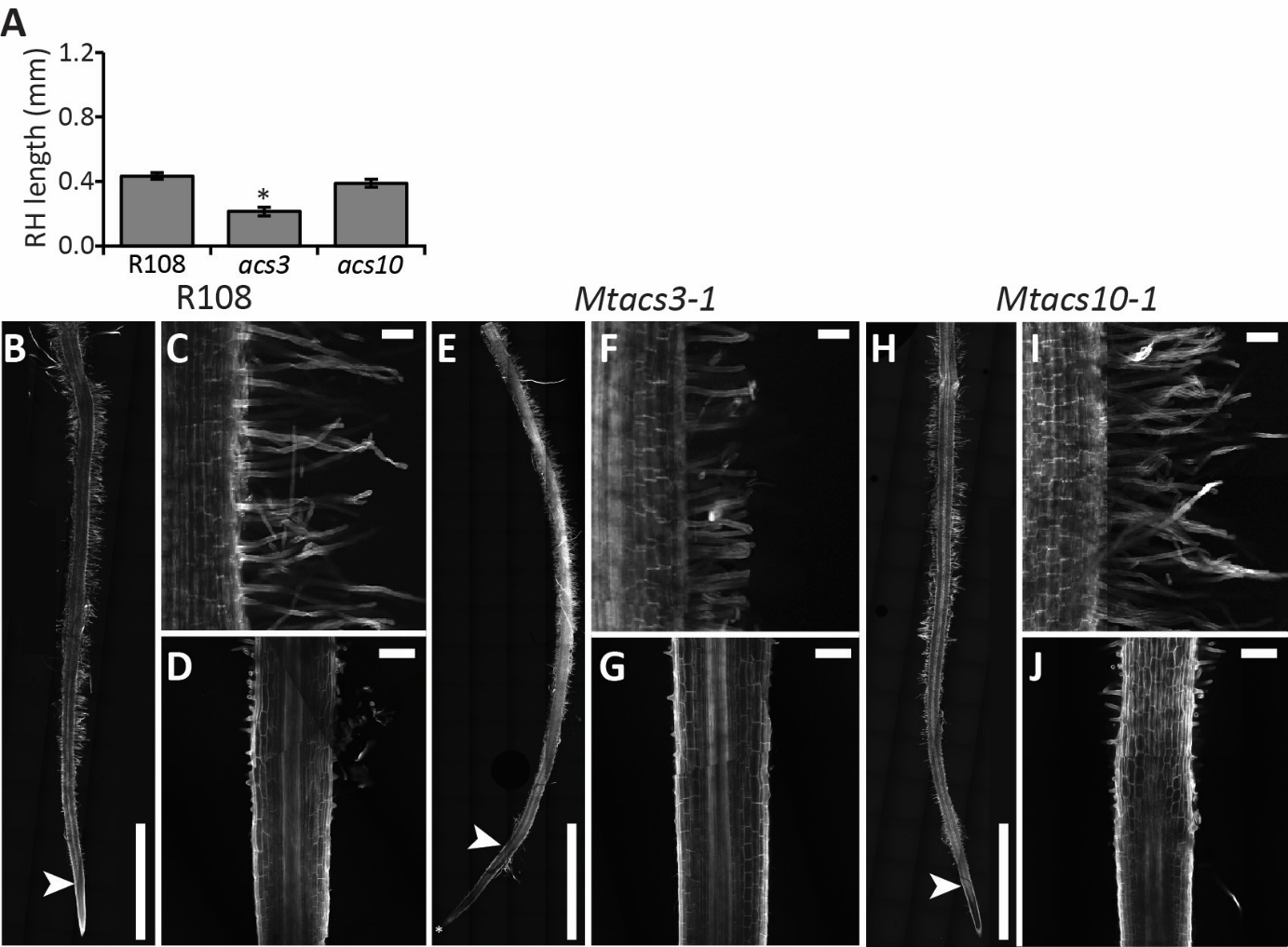
**Supplemental Figure S13:** Root hair development on wild-type Medicago R108, *Mtacs3-1*, and *Mtacs10-1*. **(A)** Average length of the root hairs (RH) in the susceptible zone measured at roughly 4 mm distance from the root tip in wild-type R108, *Mtacs3-1*, and *Mtacs10-1.* Bars represent mean ± SE; n>6 (multiple root hairs from minimally 6 plants measured); an asterisk (*) indicate significant differences (paired Student t-test, *P* < 0.05). **(B-J)** Representative confocal images of the Medicago **(B-D)** wild-type R108, **(E-G)** *Mtacs3-1*, and **(H-J)** *Mtacs10-1* root tips and root hair zones. **(B, E, H)** Confocal tile image of the Medicago **(B)** wild-type R108, **(E)** *Mtacs3-1*, and **(H)** *Mtacs10-1* root (Arrowhead points at first root hair, asterisk (*) in E marks root tip, scale bars 5 mm). **(C, F, I)** Confocal image of root hairs in the root susceptible zone of Medicago **(C)** wild-type R108, **(F)** *Mtacs3-1*, and **(I)** *Mtacs10-1* (scale bars 100 µm). (D, G, J) Confocal zoom in image of first root hairs of Medicago (D) wild-type R108, (G) *Mtacs3-1*, and (J) *Mtacs10-1* (scale bars 200 µm).

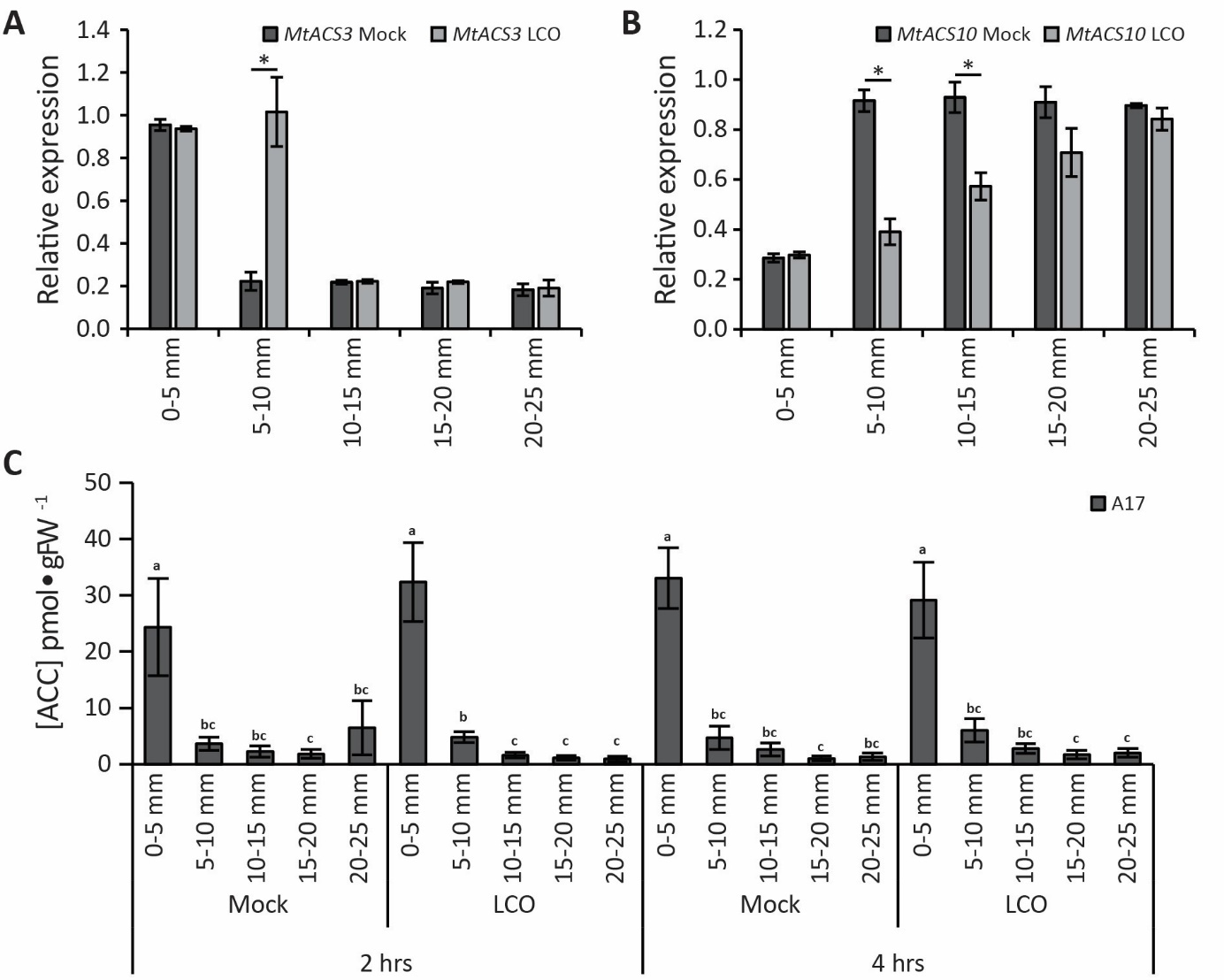
**Supplemental Figure S14:** *MtACS3* and *MtACS10* expression dynamics and ACC concentrations along the Medicago wild-type Jemalong A17 root axis and R108 mutants. (A) Relative expression over the root axis per 5 mm zones under Mock and lipo-chitooligosaccharide (LCO) application in Medicago A17 of (A) *MtACS3* and (B) *MtACS10*. Bars represent mean ± SE; n=3, independent biological replicates each consisting of the susceptible zones of ~16 pooled plate grown plants. (C) Concentrations of ACC along the root axis of Medicago A17 per gram fresh weight (FW) in response to Mock or LCO treatment (2 and 4 hours). Bars represent mean ± SE; n=5, independent biological replicates each consisting of ~40 pooled mass inoculation plate grown root tips separated into 5 segments (0-5 mm, 5-10 mm, 10-15 mm, 15-20 mm and 20-25 mm measured from the root apex); different letters indicate statistical significance (one-way ANOVA followed by Tukey’s HSD test, *P* < 0.05).

**Supplemental Table S1.** Overview of ethylene biosynthesis genes with version ID and common name used in different publications. Based on Gómez-Fernández et al., (2025). “-“ gene absent from the dataset, “nn” gene was not named with a number.

| Common Gene Name | A17  (MtrunA17r5.0-ANR) | A17  (Mt4.0v1) | Breakspear et al. 2014 | Larrainzar et al. 2015 | Van Zeijl et al. 2015 | Schiessl et al. 2019 |
| --- | --- | --- | --- | --- | --- | --- |
| ACS1a | MtrunA17_Chr8g0389311 | Medtr8g101750 | - | - | **ACS1** | *nn* |
| ACS1b | MtrunA17_Chr8g0389321 | Medtr8g101820 | - | **ACS7** | - | - |
| ACS2 | MtrunA17_Chr4g0054371 | Medtr4g097540 | - | **ACS4**  contig_12436_1 | **ACS2** | *nn* |
| **ACS3** | MtrunA17_Chr6g0488041 | Medtr6g091760 | Mtr.20234.1.S1_at | *nn* | **ACS3** | *nn* |
| ACS4 | MtrunA17_Chr3g0135251 | Medtr3g103550 | - | - | - | - |
| ACS5 | MtrunA17_Chr5g0398431 | Medtr5g011400 | - | - | - | *nn* |
| ACS6 | MtrunA17_Chr5g0400831 | Medtr5g015020 | - | **ACS8** | - | *nn* |
| ACS7 | MtrunA17_Chr6g0475681 | Medtr6g463260 | - | - | - | *nn* |
| ACS8 | MtrunA17_Chr7g0248681 | Medtr7g079080 | - | *nn* | - | *nn* |
| ACS9 | MtrunA17_Chr8g0348351 | Medtr8g028600 | - | - | - | *nn* |
| **ACS10** | MtrunA17_Chr8g0387481 | Medtr8g098930 | - | contig_49261_1 | - | *nn* |
| ACO1 | MtrunA17_Chr2g0289341 | Medtr2g025120 | - | *nn* | - | *nn* |
| ACO2 | MtrunA17_Chr3g0121141 | Medtr3g083370 | - | *nn* | - | *nn* |
| ACO3 | MtrunA17_Chr3g0125201 | Medtr3g088565 | - | - | - | *nn* |
| ACO4 | MtrunA17_Chr5g0439011 | Medtr5g085330 | - | *nn* | - | *nn* |
| ACO5 | MtrunA17_Chr6g0488511 | Medtr6g092620 | - | - | - | *nn* |
| ACD | MtrunA17_Chr8g0393311 | Medtr8g107670 | - | - | - | *nn* |

**Supplemental table S2.** Primers and Gene IDs (Medicago genome v4.01) used in this study.

| **Primer name** |  | **Sequence** |  | **Gene ID Mt4** |
| --- | --- | --- | --- | --- |
| MtACO1_Fw |  | ATTGGAGAAACTGGCAGAGG |  | Medtr2g025120 |
| MtACO1_Rv |  | CCAGCATCTGTGTGTGCTCT |  |  |
| MtACO2_Fw |  | CAAAGGGACCAACTTTTGGA |  | Medtr3g083370 |
| MtACO2_Rv |  | GGGAGGGACATCTACCCAGT |  |  |
| MtACO3_Fw |  | CCGGAACTTGTGAATGGTCT |  | Medtr3g088565 |
| MtACO3_Rv |  | ATCTGCCATTGCTCAGGACT |  |  |
| MtACO4_Fw |  | CATCGTTGTCAACCTTGCTG |  | Medtr5g085330 |
| MtACO4_Rv |  | GGAGCAGGGTAAATGACAGC |  |  |
| MtACO5_Fw |  | GATGCTGGTGGAATCATCCT |  | Medtr6g092620 |
| MtACO5_Rv |  | TCCATTCTTGTCAGGCATCA |  |  |
| MtACS1_Fw |  | CGGAGATGCTTTGCTTGTTCC |  | Medtr8g101750 |
| MtACS1_Rv |  | CAGCACTCCACTCACTTTCATG |  |  |
| MtACS2_Fw |  | TGGTTTCGTGTGTGTTTCGC |  | Medtr4g097540 |
| MtACS2_Rv |  | TTGCCAGTGGACTCAACAAC |  |  |
| MtACS3_Fw |  | TGGGCTTGGCAGAAAATCAAG |  | Medtr6g091760 |
| MtACS3_Rv |  | ACCACCTTTTCCCCATGTTG |  |  |
| MtACS4_Fw |  | TGCAACTTTTGAGGCAGAGA |  | Medtr3g103550 |
| MtACS4_Rv |  | AATCCTTCGCATTGCAACTT |  |  |
| MtACS5_Fw |  | CTGCTGGGCTGACATGAGTA |  | Medtr5g011400 |
| MtACS5_Rv |  | CCCGATCCGTTCCACTACTA |  |  |
| MtACS6_Fw |  | TCATGTTTTGCCTTGCTGAA |  | Medtr5g015020 |
| MtACS6_Rv |  | AAGCCGATTCTGTGATTTGG |  |  |
| MtACS7_Fw |  | CAACGGTTTTCTCCTGGGTA |  | Medtr6g463260 |
| MtACS7_Rv |  | CGAAATAGAGCCACGAGAGG |  |  |
| MtACS8_Fw |  | GGGTCTTCCGGGTTTTAGAG |  | Medtr7g079080 |
| MtACS8_Rv |  | AAAATTCACGTCGCTCTGGT |  |  |
| MtACS9_Fw |  | ATCCAGGATTCGTTCGTGAC |  | Medtr8g028600 |
| MtACS9_Rv |  | CTAATGGGTTGGAGGGGTTT |  |  |
| MtACS10_Fw |  | CGAACTCGTCATCAGACAGC |  | Medtr8g098930 |
| MtACS10_Rv |  | TGACCGTAACCTCGTTCACA |  |  |
| MtACD_Fw |  | AAACAAAGTGCGGAAATTGG |  | Medtr8g107670 |
| MtACD_Rv |  | GGGATCTTGGTCAACGAGAA |  |  |
| MtNIN_Fw |  | GGGAGAAAGTCCGGGGACAA |  | Medtr5g099060 |
| MtNIN_Rv |  | GACACACACCGATGCTCTTTGC |  |  |
| MtACT_Fw |  | GCAAAGGCAGAATATGATGAAT |  | Medtr2g008050 |
| MtACT_Rv |  | CCACTATGACTGCCAGAACACTTA |  |  |
| MtPTB_Fw |  | TGAACCAGTGCCTGGAATCCT |  | Medtr3g090960 |
| MtPTB_Rv |  | CGCCTTGTCAGCATTGATGTC |  |  |
| MtUBQ_Fw |  | CACCTCCAATGTAATGGTCTTTCC |  | Medtr4g091580 |
| MtUBQ_Rv |  | CCCTTCATCTTGTCCTTCGTCTG |  |  |
| MtACS3_RNAi_Fw |  | CACCCCCTACTTTGCTGGATGGAA |  | Medtr3g088565 |
| MtACS3_RNAi_Rv |  | CAACTGCGGCTAATGAGCTT |  |  |
| ACS10_RNAi_Fw |  | CACCAACCACCCCATACTTCATTCC |  | Medtr8g098930 |
| ACS10_RNAi_Rv |  | CCAACCAAAAATCCTCAAGG |  |  |

**Supplementary Table S3:** Gene probe sets used for ViewRNA *in situ* hybridization

| **Gene name** | **Gene ID Mt4 / r5.0** | **Assay ID according to ThermoFisher Scientific** |
| --- | --- | --- |
| *MtACS3* | Medtr6g091760 / MtrunA17_Chr6g0488041 | catalogue number VF1-6000770 |
| *MtACS10* | Medtr8g098930 / MtrunA17_Chr8g0387481 | catalogue number VF1-6000771 |

**Supplemental table S4.** Sequences use for RNAi constructs.

*MtACS3^i^* _fragment
^CACC^CCCTACTTTGCTGGATGGAAAGCATATGATGAAAACCCTTATCATGAATTAACTAACTCTTCTGGTGTTATACAAATGGGATTGGCAGAAAATCAAGTTTCATTTGATTTGGTAGAAAAGTATTTGGAAGTGCACCCGGAAGATTACAATGGTTTCAGAGAAAATGCATTATTTCAAGACTATCATGGACTTAAATCATTCAGAACTGCAATGGCAAGTTTCATGGAACAAATAAGAGGTGGTAAAGCTACATTTGATTCGGAAAGAATAGTCATCACTGCCGGAGCAACTGCGGCTAATGAGCTT

^Note: CACC added for TOPO cloning^

*MtACS10^i^*_fragment
^CACC^AACCACCCCATACTTCATTCCTACACATTGCAATCTCTTCTAACTTATTCTTACACCTTCAAACATAAGAAGAATTCCCACACAAATTCGCTTCTTCCTCGTACTTTTTTGTTTTTGTTTTTGAATCTCATTGATTTCATCACATTTTCTATTAAATTAATTAATATTTTAGACTATAGTAATTAATAATGGGACTTGTGAGCATGGACCAACCTCAATTGTTGTCCAAGATAGCCACTGGTGATGGACATGGTGAAACATCATCTTACTTTGATGGATGGAAAGCTTATGATAAAAACCCTTTTCATCCAACCAAAAATCCTCAAGG

^Note: CACC added for TOPO cloning^

**Supplemental table S4.** Sequences used for ectopic expression of *MtACS3* and *MtACS10*.

EC74831; *MtCRE1p-MtACS3-t35S*

EC74832; *MtCRE1p-MtACS10-t35S*

EC74833; *MtCRE1p-GUS-t35S*

EC74963; *MtACS3p-GUS-ACS3-3`UTR*

EC74964; *MtACS10p-GUS-ACS10-3`UTR*

*MtACS3* ATGGGTCTTGAGATTGAACAAGAACACCCTTGTGTTGAACTTTCAAATATTGCAACTTCTGAAACTCATGGAGAAAATTCTCCATACTTTGCTGGATGGAAAGCCTATGATGAAAACCCTTATCATGAAATAACTAACCCTTCTGGAGTTATACAAATGGGCTTGGCAGAAAATCAAGTATCATTTGATTTACTTGAAAAATACTTGGAAGAACACTCAGAGGCTTCAACATGGGGAAAAGGTGGTTCAAGTTTTAGAGATAATGCATTATTTCAAGACTATCATGGACTTAAATCATTCAGAAAAGCAATGGCAAGTTTCATGGAAAAAATTAGAGGAAATAAAGCAAAATTTGATTATGAAAGAATCGTCCTCACTGCTGGTGCTACTGCTGCCAATGAGCTCTTGACTTTCATTCTTGCAAATCCAGGAGATGCTTTACTTGTTCCAACACCATACTATCCTGGATTTGATAGAGATTTGAGATGGAGAACTGGTGTAAACATAATTCCAATCCATTGTGATAGCTCAAACAATTTTCAAATCACACTTGAAGCATTAGAAACTGCATACAAAAATGCAGAATCCATGAACATGAAAGTAAAAGCAGTACTTATAACCAACCCATCAAATCCATTAGGCATATCGATTCAACGTTCAGTTCTCGAGGACATTCTGAACTTCGTGACTCGCAAGAACATACACCTTGTCTCAGACGAAATCTACTCGGGCTCAGTTTTCTCTTCACATGAATTCATAAGCATAGCCGAGATTCTTGAATCTCGTCAATACAAAGACGCGGAAAGATGTCACATTGTTTATAGTCTTTCTAAAGATCTCGGTCTACCAGGTTTCAGAGTCGGAACAATTTATTCCTACAACGATAAAGTTGTTACAACAGCACGAAGAATGTCGAGTTTTACCTTAATATCTTCACAAACACAACATCTTTTAGCATCAATGTTGTCAGATGAAAGTTTCACTGATAATTACATCAAGGTCAATAGAGAAAGATTAAGGAAAAGATATGAAATGATCATTGAAGGTTTGAAAAGTGCTGGAATTGAATGCTTGAAAGGTAATGCAGGGTTGTTTTGTTGGATGAATATGAGTCCAATGTTGGAAAGTAATACAAGAGAAGGTGAATTGAAGCTTTGGAATGAGATTTTGAATGAAGTTAAGCTTAATATTTCACCAGGGTGTTCTTGTCATTGTTCCGAACCCGGTTGGTTTAGGGTTTGTTTTGCAAATATGAGTGAAGAAACACTTGAACTTGCACTCAAAAGAATACGTGATTTCATGAATAACAAGGACAGAAAGGATATAGGAATATAA

*MtACS10* ATGGGACTTGTGAGCATGGACCAACCTCAATTGTTGTCCAAGATAGCCACTGGTGATGGACATGGTGAAACATCATCTTACTTTGATGGATGGAAAGCTTATGATAAAAACCCTTTTCATCCAACCAAAAATCCTCAAGGTGTTATCCAAATGGGTCTTGCAGAGAATCAGCTTACTGCTGATTTGGTTCAAAATTGGATAATGAGTAACCCAGAAGCCTCAATTTGTACTCTAGAAGGAGTACACAATTTCAAAGAAATGGCTAATTTTCAGGATTATCATGGTCTACCAGAGTTCAGAAATGCTGTGGCTAAATTCATGTCAAGAACAAGAGGAAATAGAGTGACATTTGATCCTGATCGTATTGTCATGAGTGGTGGAGCAACTGGAGCACATGAGGCCACTGCCTTTTGTTTGGCAGATCCTGGTGATGCTTTTTTGGTGCCTACACCTTACTATCCAGGATTTGATCGAGATTTGAGGTGGAGAACAGGGGTTAAACTTGTTCCAGTTATCTGCGAAAGTGCAAACAATTTCAAATTAACAAAACAAGCCTTAGAAGAAGCATATGAAAAAGCCACAGAAGATAACATCAGAATTAAAGGTTTACTCATAACAAATCCCTCAAATCCATTAGGCACAGTTATGGACAGAAACACATTAAGAACCGTTGTAAATTTCATCAACGAAAAGCGTATTCACTTAATAAGCGATGAAATTTACGCTGCAACGGTTTTTAGCCACCCAAGTTTCATAAGCATAGCTGAAATATTAGAACATGACACAGACATTGAATGTGACCGTAACCTCGTTCACATAGTTTACAGTCTTTCAAAAGACATGGGATTCCCTGGTTTTAGAGTTGGTATAATATACTCTTATAATGATACCGTTGTAAATTGTGCACGAAAAATGTCAAGTTTTGGATTAGTTTCAACACAGACACAATACTTGATGGCGAAAATGCTGTCTGATGACGAGTTCGTTAAAAAGTTTCTTACTGAAAGTGCAAAGAGGTTAGCACAAAGGTACAGAATTTTCACCAGTGGATTAACCAAAGTTGGAATTAATTGTTTACAAAGTAACGGTGGACTTTTTGTGTGGATGGATTTGAGAGGACTTCTTAAGGAAGCTACATTTGAATCAGAATTGGAACTATGGAGAGTGATTATTCACGAAGTTAAGATTAATGTTTCACCTGGAGTTTCTTTTCATTGTTCTGAGCCAGGGTGGTTTAGAGTGTGTTATGCTAACATGGATGATAGAGATGTGCAAATTGCTTTACAAAGGATTAGGTCATTTGTGGTTCAGAATAATAAGGAGGTTATGGTGTCTGAGAAGAACACTAAACCTTGTTGGCATAGTAATTTGAGGTTAAGCCTTAAAACAAGAAGGTTTGATGATATTGTAATGTCACCTCATTCTCCATTTCCTCAGTCACCTCTTGTTAAAGCCACTACTTGA

*GUS* (pICH75111) ATGGGTCAGTCCCTTATGTTACGTCCTGTAGAAACCCCAACCCGTGAAATCAAAAAACTCGACGGCCTGTGGGCATTCAGTCTGGATCGCGAAAACTGTGGAATTGATCAGCGTTGGTGGGAAAGCGCGTTACAAGAAAGCCGGGCAATTGCTGTGCCAGGCAGTTTTAACGATCAGTTCGCCGATGCAGATATTCGTAATTATGCGGGCAACGTCTGGTATCAGCGCGAAGTCTTTATACCGAAAGGTAAGTCTTACTCTCTCTTTTTTGGTCTGTATTTTTAATTTTTTGAAGTATACTATTTGTACTGACGCTAATAATCTTTTTTCAGGTTGGGCAGGCCAGCGTATCGTGCTGCGTTTCGATGCGGTCACTCATTACGGCAAAGTGTGGGTCAATAATCAGGAAGTGATGGAGCATCAGGGCGGCTATACGCCATTTGAAGCCGATGTCACGCCGTATGTTATTGCCGGGAAAAGTGTACGTATCACCGTTTGTGTGAACAACGAACTGAACTGGCAGACTATCCCGCCGGGAATGGTGATTACCGACGAAAACGGCAAGAAAAAGCAGTCTTACTTCCATGATTTCTTTAACTATGCCGGAATCCATCGCAGCGTAATGCTCTACACCACGCCGAACACCTGGGTGGACGATATCACCGTGGTGACGCATGTCGCGCAAGACTGTAACCACGCGTCTGTTGACTGGCAGGTACTTCATGCTTCAACGTGTAACTTAAGAGATACTGTGTGAAATTTTATATTTCCATACATTTGCTTGACCTTTGCTTTTTGTCAATTTTTTTCCCCTTACAGGTGGTGGCCAATGGTGATGTCAGCGTTGAACTGCGTGATGCGGATCAACAGGTGGTTGCAACTGGACAAGGCACTAGCGGGACTTTGCAAGTGGTGAATCCGCACCTCTGGCAACCGGGTGAAGGTTATCTCTATGAACTGTGCGTCACAGCCAAAAGCCAGACAGAGTGTGATATCTACCCGCTTCGCGTCGGCATCCGGTCAGTGGCAGTGAAGGGCGAACAGTTCCTGATTAACCACAAACCGTTCTACTTTACTGGCTTTGGTCGTCATGAAGATGCGGACTTGCGTGGCAAAGGATTCGATAACGTGCTGATGGTGCACGACCACGCATTAATGGACTGGATTGGGGCCAACTCCTACCGTACCTCGCATTACCCTTACGCTGAAGAGATGCTCGACTGGGCAGATGAACATGGCATCGTGGTGATTGATGAAACTGCTGCTGTCGGCTTTAACCTCTCTTTAGGCATTGGTTTCGAAGCGGGCAACAAGCCGAAAGAACTGTACAGCGAAGAGGCAGTCAACGGGGAAACTCAGCAAGCGCACTTACAGGCGATTAAAGAGCTGATAGCGCGTGACAAAAACCACCCAAGCGTGGTGATGTGGAGTATTGCCAACGAACCGGATACCCGTCCGCAAGGTGCACGGGAATATTTCGCGCCACTGGCGGAAGCAACGCGTAAACTCGACCCGACGCGTCCGATCACCTGCGTCAATGTAATGTTCTGCGACGCTCACACCGATACCATCAGCGATCTCTTTGATGTGCTGTGCCTGAACCGTTATTACGGATGGTATGTCCAAAGCGGCGATTTGGAAACGGCAGAGAAGGTACTGGAAAAAGAACTTCTGGCCTGGCAGGAGAAACTGCATCAGCCGATTATCATCACCGAATACGGCGTGGATACGTTAGCCGGGCTGCACTCAATGTACACCGACATGTGGAGTGAAGAGTATCAGTGTGCATGGCTGGATATGTATCACCGCGTCTTTGATCGCGTCAGCGCCGTCGTCGGTGAACAGGTATGGAATTTCGCCGATTTTGCGACCTCGCAAGGCATATTGCGCGTTGGCGGTAACAAGAAAGGGATCTTCACTCGCGACCGCAAACCGAAGTCGGCGGCTTTTCTGCTGCAAAAACGCTGGACTGGCATGAACTTCGGTGAAAAACCGCAGCAGGGAGGCAAACAATGA

*MtCRE1_pro_* GGAGCCTAGAACCAATATAAAGACTATTTTTATTGTCAAAATCTAGACTTATTAGAACTTTTTTGTTTTCTAATTTCCAAATGAGTTATATTATATGAACAATTTTTTCTTGTGACAACTTGATTGACAATCAAATTATACAAAGAAATTTAGATAAATATACAAAAATCAAAGCAATAGAGAGAGAAAGTAGATAAATAATGTGAGTATGAGAGATAAAATTGTCACAAAAGTTGTCAAAAATGATTGTTCAATTATCATTTTTCTTTTCAAATATTGAACTTTTTCTTACAAGATACAAATGAGGCATTCTTCGATAGTCACGAACACAAATGCATAGTGTTTGGACACACACATATAATTTATATGTCATTTCTTGAAATACTAAATTATTTTATTATCAATTTCTTTTTCATTATTAAAAAAAATTGTAATGTGTACGCGCCTAAACACTATCCATTTGTATTTGTGATTTCTCGATGCCAAGACTAAACTTGATGGCAATTATTTTATTGAAAAATTTGATAGCAATTAGTACCACTTAAGAATTAAATTGGTGGTATATTCATCTATTCTTTCTTTAGAGATGTAATCATGCCTTTATTTGAATAAAGTGCCACAACGAGATAAGGTCACCTTAATAATTTGGAAGAGGACATTCTATCTTTCTTTTTTTAACATTACATTTATTATTTTTAATGAAGTCGCAAAGATGTAAAAAAATTGTCCAAACAACAAAGTAGGGAACCAATCAAACCGTTTTTTCTTTTTCGTTGCAAAGTTAAGCTAAATGAAATGAGTAGCAATATTTGAATAACCATGTTGGATAACTTATGTGATAATTTTTTTTCTCTCTTTTTATTGGTCAAAATCAATGGAAAGAGAAAAATAAGAGAGAGAAAAAAAGTATAATATGAGTATGAGAGAGAAATTTGTAAAAAAAAAAAATCATAAAATGGTTATACAAATGTAATTTATCAAAAGAAATAACTAAATTAACTGTTTACCAACAATGAATGTATGAGTACTACTATACATAGATTATTAAGGATACATTAGTCACAACATTAATTAATACAAAACAACAAAGTTAATTAATGTTGATTATCCATTGAGGGTAGTTATGGAACTCTAAATAATAAAAAGGGTGGGTGTTTAGGGTAATTCACATGGGAATAAGATCCTAGCATTGTCGCAAGTCACTATCTTTCAGATTTGTGATTGTGATTTTTTCTATCTTTCTCTCTCCTCTGTTCCACTACACAACATTGTTTAACGTTGGAACACATAGAAATAGTGAGAAAGACCCATTTGAAGATTCAACTCTAGAATGTGAAAAAGTTTCAATTCTTACATTCATTTTCCAAAGTTAGTAATAATAGCTTAACTGGGTCAGTTCATTCTCCATTGAAGCTTCTTTTCAAAGTGCTAGTTGAAAAAAGATGCTAACTTGAGGAGAAAATATTGTTGGGTTGAAAGAGAGAGTTTTGACTTGACTAGAGTGTGAAGTAGAAGGGAGACAACAACAGCGTATCAGAGAAAATCGAAAAAAATTCGATTTAGTTTTAGTAATGTCCTGTTTTGTGCTTAAAAAATAAAGAAAGTAAAGTAAGAAAGCATATATAAAAAGGTTGTATAAAAAACAAAAACTAGTACAGAAAGAGAAAGGTGCAGTGCAGTGCAGTGCAGATCAAGACGAACCCAACCAACCAATCTGCTCTCAACTGTGTTCATTGTATTGTTTCTCTTTCCTATTCCTATACAGGTAAACATTGTTTCTCTAACTTACTCATTGGGTTCTGTGTGTGCTTCGCTGTTTTTTTTTTTTTTTTTTGTTTAGTATGGTTTGTGTGTTTTATTTTTGAGGTCAATGTTTGGTTTTTTTGATTAAACAAATAAAAGTGTGAACGTGCTTTTGTTTGTCCTTGAGCAGAGACTGGTTTCACGCCTAGAGAGCATGATTAGAAGTCTCTTGAATTTTTAATTCATAATATTGGCTGTTGAGAGAAAGGAGAAGGTGCTTCTTTGGAATTTGAGCTGTTTTTCCATCTTTTGAGAGCCATGCTTGTCTCTATTTCCCTTCTTTTTAATTCTTTCTCTCTTTTCTCTCACTTCTTTTTTTTATTCTCTAATTTTCTGTGTAATTTTTCAAATGAATTTTCCTCAATCATTCATCATGCTTGGTTTTGCTTTTTTTATTTGATTCTTGTTTCCTAAGAGGCTTTGTTTTGGTTTGGTTTGAATCACGCAAAACAATGCTGTAATGATGCTGTACTTGGTTTTTTGTTCTGTGAACTTTTTCTTTCAACCATCAGATAAAAGTGTCTGAGATTTTAGGACTACTTTACACTTTAAGCAAAGTAAAAGCTACTTCCTATATATAAAAAAGTTTGTGCTCTGTTAATTGAGTATCTTTTGGGACTTCCATAGATATGAAGAAGAGTTCAGAGAAATAGGCAGAAAGAAAGAGATTTGGTGTTAGTTGGTGTTGTAAATG

*t35S* (pICH41414) CTTCTCTAGCTAGAGTCGATCGACAAGCTCGAGTTTCTCCATAATAATGTGTGAGTAGTTCCCAGATAAGGGAATTAGGGTTCCTATAGGGTTTCGCTCATGTGTTGAGCATATAAGAAACCCTTAGTATGTATTTGTATTTGTAAAATACTTCTATCAATAAAATTTCTAATTCCTAAAACCAAAATCCAGTACTAAAATCCAGAT

*MtACS3p*

AACCCCCTAATTTCCTAACAATAACAAATTGTTTGGTTCAGTTTTAACGTTAAAAATATGAATTGGACACTCAACTCAAACCAAAGACAATTGATCAGATTCAGTAATGAGTTTGGTCAAATTAGCCCAACCCAACCTCCAAACACTCCTAATCAGAGGTGGATAAAGTGGTGGTCGTCAATGGTGGGTGGTGGTTAGGGTTTACTGGGGGTAACATGAGTGGTGGTAGAAGGTGGTCAAAATAATGGTCAGATAGGTGATAAGAGTTGGCCAACAGTGGTCAACAACCGACCACTGGACGGTGGTACGCGTTGTGGACTCTATAATACTTCAATATATTTAATACTATAAAAAAAAAAAAAATTTAATATATTTAAGAGATTTTTTTTGGTCAAGTAGACTAGTGGTTAGAATTCCCCTTTTTTCAAAGCGAATAAGTGGGGTGTCCGGGTTCAAATCCGGACCCCTGCATATAATAATGCATATCTTTTCCAACTGAGCTATGCTCACCGGAATTATATTTAAGAGATTTATTGTCGTAGTATGTTTATTTTTTTCATTTTTGAAAGGAGGCATTGTTATAGTTTAAAAAATCATAAAAAAGAAAAAAAATTGATTACGTTAGAAACTTTACAAAAGACTGATGTAATCCAAATTAGTTTTGCCAAAATACTAAAAGTGTACATTTTCTTCTTCTAAAAGTTTGTCTTGTCACAAAAGAAAAAAGTTTGACATAGAAAATGAGAATTCATTTTTTTTAGTATCATTTCTAATAATTAAATTCAACTGAAACAATTTCCAAGCCTATATTCGTTTTGAAATTCTTCGTCTTTCGTCAGATTTATTACTTTTTTCTTTTGTCGTTGCTCATTAGTTTATTTATTTATTATTTTTCATGTTGACTTTTGCATGGAATACAACTTCTTCAATTTCTCAAATTGAAAGTTTGTGTGAACCTGTGAATTATGTGCAAAGATGATTAAGCATTTCACTCATATTAGTATACTTATTAATTAATTTATACTTAATGAATAAGTATACGTAAATTCTCAATAAGGTTGTGGTTCTCTTAATTTTTTAATATTAAAAGAAAAATAGCTTTCATTTTCATAATGATTTTTCTTAAAGGATTCATTTTCTTAATGATGTTATTTTGCTTTTAATATAAAGCTCTCATTGAGCCACGGTGTATATATATATATTCCTAATAAGTAAACGTCCCATAAATGATATTTGTACAACCATTTTGTAATAACTTTTGTGACAACTTTCTCTCTCATACTCACATATGTTTTCATTTTATCTTTCTATTGCTTTGATATTTGTGCTAAAACTTATTTTTTCTTTATAAATTTATGGTTGTCAAATAAATTGTCTATCAAAAGGTTATTCAAATAACACACCTCTTTTATCAATATACACCTCATCCTTAAAAAATAAAATCAATATACACCTCATAAAAAAGAACTACAATATGCCAAAAATAGCAAAAGTTAAACCATCCCTTTAAAGAATTTTACTCTCCCAAGTCTTAACACTAGTATTAATCAAGCAGACAGCTATAATTACATATGTTTAGCTGTTTGTACCAAAAATATTAATATTGTTGGTTTTGAATTCCGGATAAATGTTCTGAGTCATAGGTATGCCCTTAACCCTAATGAATCCCGGAGCCCAACCCATATAAACTTTATCTTATAAAAAAATAACTCTAGAAGTATATGAAAATTTTGTGATATTTTTCATTTAACAAGACAATATTATTTAAGGCCAATGATAAAACATATAATTTTGAAATGATGAATTATGTGATATTTTCCATAGATAACTAATCTATTATTGAAATAATTGTAAATTATGATTATTGTTTGTGAGATGCCTATCATTTAGTTTTATTGATCGCCTATAATTTTTTCCGATGATAGTTAAGTCTAGTAGAATGAGTTAGTCTCCTACAATTTCGCAAACCACAATTTGATTACTCATTATATATAGTGGCACCCATATTTAGTGTCCTATTTCTACTCAAATAAGATTGCCTCTAGATTGCCTTTCATTTCATCACCAAGGAAAAAATGGTGCTCCAAATATTTAGGAAGTGCACAAGTAAATGACTAATTAGAACAAGGTGCACCACAATTAGATTAGATACAACTACAAACTTTAAAGGTTGTATATATTGAAGAGATTTGATGCATAAACCCTAAAAATATATTTGATTATACCCAAAAAAAAAATCATATAGAAATTGATTGTATTGGATTGGATTGGAGTACTCATTGTGCTCCTTTATCCAATAATCTTAATTTGTTCTTCAAATTAAGGGAGATGTTTTATTTAAATCTCACAAAACTCATTGATCAATTTTGTTGTTTGGTATTGATGCAAAAGTGGGGTCACAAGATATCCCAAGATGTTCTCTTTTGTCATAGTTCTTCTTGCAAAATCAGAATCTTAATCTATCTTTTGGGTTACATGGTATCTTTGTCTGGTCATCAAAACTTAACTACTCCTTGCTAGAATTTTCTCCATTTTTTTCTTATATGCTAAACACCCAATTCAGATCTTAAATCATGATTTGGTTTCCACAATGGAATTTTTGGTGTTTATTTGGCATTGGATTTGATTAGTCGTAGGTTGAGAATAGTAGAATTGTTTTTTCAAGAACTCAACCACCTGTAAATTTAATAAACAAGTATGAAACCATAAATTAATTAAGCCACAAACAAAGTTATGAACAATCTGCTCCATTAAAACAAAGGTTATTTTATTTTTTTATAAAATGTTGAAGTAAGTTGAACCCGAAGAGTTGCGAGGATCAAGAAAAATTTCAAGATTCAGAGACGACAAATACAAATACTTACATATTATCTATTAACGATGTTCTCAAACTAACAACGAACATCAAACTCAGGTCTTTGAAGAATCTGAGCGGAATCCTTATCAACGAAGATCAAGACAAATTTCAAGATTCATAGACCACAGATACAAGTACTTACATATCATCTAATAACGATGTTCTTAAACTAACAATGAACATCAAACCCAGATCTTTGAAGAATCTGAGCGGAATCCTTATCAACTTCACCAAACAAAGTTGGTAATTTGTAAGCAAAATTAGGGAACTATAATAGCAACTTAATCAAAACATCCATCTCCAAACAACCACCATATAATTTCCGTGTGGATCTGTCACAGTTGTCACAAGTAGCAACAGATAAACTATGTGAAATTAAATTAAATTAAAGCTTAAAAGGTAAAGCTTTGATTTGCTTACACTACAAAATCAATAATTTATATTATATACATTTTATGCATTATTTATGCTGAATAAGCCAGCCAAGTTGAAAAGGGACCTTTTTTCCTTGTCTTTAAGCCTCAAAGCCTATATATAGAGAAATTTCACACAAACTTTTCATTCAACTCTCTCTCACCCATTTTCTCTCTCTACCTATCTAACTACCTTCTCTCTCTTTCCCCTCCTTCTAAAAATAGTTAGT

*MtACS3-3`UTR*

TTTTTACCATCACTCTACCATGATTAAATAGTATTTTTTTAGGGTCAAATCAATCAGAAGAGTCAGAGTTTGAATCATGACAAAAATAATCATTTGTCAAGTTTTACTTATCTCCCAAACGAATTCCGATTACCAGTTTCTTTCGCCTTGAAATGGGAGGGATAAGACCAAAACATTAGAGTCTTTGTTTATTTATTTTTATTTTTTTTCAAATGGTTGAGGCCATTTTGGTATATTCTTTTAAAAGCACATAATGTAAAGGTGAATATTCAATTGAAGTGTATCAATCCCTTGTTTGCTTGCTTAATTTACTTAAGCAAGTTTTTATTACAAAATTAATCTTTTGTATTATTTTGTTTTATAATGAATGAAATTTGTTACTATTAAATTATGTGTCTTAATTACTACATTATTTGTATGATGAGTTAGTATAGTGTCTATATCATCTTGAACATCTTGATTGGTTGTGTAGTAAAAAATTAATCCTTATTGAATGAGTATTGGTCTTGAATTTATCAACATCATTTGACTTAGTAATTACTGATTTAATTTCAATTCATATCATAGTTGACTTAGAGTTTGGAGTTTAAAGTACGTAACAGCTAGCAAAACACACCAACCAGTTATAGCTTTCAGCTTCATGAAAGGGTTGGAAACTTGGTATGAATAAAATATTAACATGCCTCAATCCCTACAGTACATGATGATGTGCATATTTTTAATTCAAGTCTAATTGCCATGTTTATTGGATAATTAGTAGTATATAAGTATGATTTGATTATAACATTGTAATTAATTATAAGTATAGTATAAGTATATACTTTGTATTAATTCCTATAGTAGTGTCGGGAGATATTCTAGAACAAATTAAAAATGGTCCACCCCACGTGATGTACACTGCTTGAATTTATATTTGTATATATAACTGCATCTCTTTGTTGTTGTTGTTGTGTTGTGTGTGTGTGAGGAAAATGATGTTGATATTGATTGATATTAAAAT

*MtACS10p*

GACCTATCGTGATTATGGTAAACACTTGAGATCCAACTCACTAAACGATAACAATATGATTTAAATTCATGTAATTGCCATCAAATTTATATGAGATTAACATGATTTAGTGAAACTGATGTGAATTAGGTTCAGCAGTGTATTGTGTGTGTGGAAAGTGGAAACCATAGTATTCAGTTCCACAATTCTGTCACCGACCAACAAGGGTAAGCAGTGCATATGGCGCCAACAACACGGCAATAATTCTATTCCAATTGTGAGTATCTTTGTTAGGTTAGTACTTAGTGGTTTGGTTACACTCTCTTTTCATTTCATTTCATGGGATGGGACATAATGCCACTTGTGTTTTCATTTAATGCTCACAGCACAAAGTCACACTCTTCTTCTTCCAACAATTCCACTCTCAATTAATACTACGTGTCAAAAATTGAAACTGTGACCCACCTTTGTAGACAGAAATAAATGAATACGTCCCATGAAAATCTATGGGTTGGTCTGGTGTAGGGTTGTACAAATCCCTCTCATGAAGGGCATCACCCTAATATAGAAAAGATATGAGGACAAAAGGTTGTCCAAGAACATTTTTTTTACGATCGAGTAAATACCAATATGGACTCTATCTTTATAAGTTATTATCAAATTAGTTTCTCATTTTATTAATTTTTTAAAATAGTCCCTCAAAAGTTTCTCAATTATATCATTGCCTCTATGTTTTTGTCATATGACGAGGAATTTAATTGACTTAATTGAAAAACTTTTGACAAAGACATGAACTATTTTTAAATATTCTCAAAAGCTTCCAATTAGATCCTTTTCTATTTGTTTTTGTCACAAGACGATAGACTTAATTGAAAAATTTTTGACAAAAATAAAAACTAATTTAAAATATTAATAGAGTCAGAGACTGATTTGAAAATTACTTACAAAGATAGAGAGCAAATTAACAGATTACTCATTTTTTTACTTATACATTTGATAATATATATTATGAAAAAAAGTCTCAAATAATTTTATCTATATTTAATTGGAGTTTTTGATATATTTAATTTTCACCAAATTAAATATAGATATTGAATCACTATCTTAACTTGCTTAAGCACTCAATCAAATTTTTCATCAGCCTTTTTGTCATGGTAGGTGGCATAATGCTTTCTTCATTCTTTCAATTTATTTTTTTCCAATACATATATATTTATGATTTAATTTGTGTAGGCCGATAATTGTAAAGCTTTTACACATTTAAATTACTCTACAATGTCCTCTTTTTAGGTTGTTCCTAAAACATCTCTACACATATCTATTTAACATTAATTTGATTAGATGCATATAAAAATGTTTTAGACTTTTTGTGCATAGAAATGAGTCTCTATATTTACATATTGATTTAAGTATTTGTCAATAAAAACTATTATATATTATTTTAGTGCGTGTTGTGTAGATGGATTACTAACTACGAAGATATCTGAATCTACATTCATATCCCCCTCATTTTTTCAAATAATTATTGGTGCTCATGCCAATATCTATAATGGTTTTTTACTCGACCAATTAATATAATTTGGTTTGCAGGTATCAAATAAATAAATAAGGCAACTTCTTCTGCGGTATACCTCACAAATTGGGGTGTACCGGTACTCCTTGCTTCATAAATTGATAAATTAACGATCATTTTAGAAAGCAAAGAGCTATTTGTCAAGGTTATTCTTTAGAAAACATCATTTCATAAGAAAAAAAAACATAATTTCATTGTTTATAAGAATCATATTAAATTTTTAACATTTTCTCATAAAAAATAATCAATAATCTTCATTATGAGTAATAAAAATTCAAAAGAATTTAATTTAATTTCATCGACATTTTGGTAAATAAGATTTAATTTTTATAATTGTCAATGATAGATTTCTTCAAAAATTCATCTAATTTAATTAATAATTTAACAATTTGGTGTATCGGTACACTCAAATATTTGGGTGTACCATATAATCCACTAGTCTCTGCCCTATAAGACACTCTCAAATAAACTATAAGATTATGAGTATTTTAAAACAATCTCTATTCTCTATATTTAACCACTTACTATTAATTTTACATTAATTAAGTCATTAAATTTTAACAATTTACTATCGATTTCCCGCAAAAAAAAAATTAATATCAATTTAATTCATAAATTTATCATTCACCATCAAATTAATCATACCATATTTTTTAAAGAACTAATATAAGTAAGATTTGATAAAGTCAAAAATATTTTTAAAATATTCATAATGCTAATTACCTATTTGAGAACGTCCTATAAAGTCTGGAATTAATTTGTTGATTTAGAGACTAATCTTCATCTCTGCACCCCTGTCCAATCCATAACTCTTTCCCATCTTTATGACCCTGGCCACATCAACTTCC

ACCAACAAATAGATAACAGATCAGAGCGTACGCAAAAATACAAATAAGTAGGTGAAACTATATGTCAATTGTATAATAAATAACAATTATACTATTTTTTTAAATAACAATTATACTAATATTATTATACTTTTTTTTTGGGTGTACATAATAATATTATTATACAAATAAAAAGTAATATTCCGTCTTATGATTATTAAAAAAAAACAGTATTATGCATAGGAACGTATACAACTGTAGTACAACACAAAACCTGAATACAAAGGTCATCACAAACTTGAAGTCTAATATTTTTCTCCCACTTTGTTGACACAAAAACAAAATACAAGTGGCATTTTGCTATTTGAACGACAATTATTACTAGTTATTGTCTGCCTACCAGCATCTGTCTAATTTTCAACTGAAGCTGCTGGCGTGGCATCATAGCTGAAAATGCTGCCAGAGTCCAAAAATAATTGCTCATTCACTATTATTAGGAGATATCGTTTAATCGATTGATTAGACTCAAATGATTAAAGAGTTTTTCTTAAAAGGAAGTGTTAGGTTTTGTTGGAGAAACTCTTTAAAGATTTAAATTTTTTTATTTTTTATAAAAGGATAAATATTTTGATTTTCAATGCATTAATTACATATATTTTCATAAAAAATTATCATTTAAAAGTATTAAAGAGTGTCCGGAGAAACTCGTCCATATTTTCCAAATAATTTTTGGATTTTGGTTTCCTATGACCCATTTTTCCTAAAAATTAAAGGCTTAACAAAAAAATATATTTGCTAGCTAGACTCTTACGCTCTCACCATTACCTTCATGTGAACCTTCCTTTTTTATATTTGGACTAACTAAAAAAATATTCTTATGTCTGAAAATCAAAATATATTTGTGCATATGCGCATGCTATAAATAACCACCCCATACTTCATTCCTACACATTGCAATCTCTTCTAACTTATTCTTACACCTTCAAACATAAGAAGAATTCCCACACAAATTCGCTTCTTCCTCGTACTTTTTTGTTTTTGTTTTTGAATCTCATTGATTTCATCACATTTTCTATTAAATTAATTAATATTTTAGACTATAGTAATTAATA

*MtACS10-3`UTR*

ATTATTTAGCATATAGAAGAAGTAATAACTGATAATTTTTTTTTTTTTTTTTTTGGTTTGTTGATTTATCAGTTTGGGAAACCACATGTTGTGTTCCTTTAATGTTGGGGGCATTTTTTCTTTTTTCACTTACTGAATCACATTGGTTTGTGGTGAAAGAAGTTCATAGAATTTTTAAGGTCAATGATGAATGAATTGTTAATAAACTTCTTCAATTATTCTACATGTTTTATTGTTGTTTTATAAACTGTTAAAGATAAATCAACTTTAGATTTGTACTAGTATGATAATGATACATATCAAATAGTATAAAGGAATTGAAGTGGATGTATGAGTAATTCACTAAAGATAACAAGACTACTGAATAAACATTCAATTTTTTATTTTTGGAATAAAGAAATTTACAAACAAGAAATTGCAGGGAGATTTACTCAAAAATATTTGTATTGAAAGTGATTCAAGTACTACCGTCCAACTTGCAAACCATCACCATGTCCACATGATTCACTTTCACGGTTTCTTTGCTATAAGATGATGCAAAGATTCAGTCTGGTCAAAAAATTGTAAAATACTATTAAAACGCAGTCTCAATTTGGATCCGTTGAATGAAAATTTCACAATTCATATTCCAAATTCATAGTTTGAATATGACTTAAGGTTGCACAAATTGAATATGAGCAGTTTCATTTGCTAAAAAAATAAGGAATCGGAATTCGGACATGATGACAACTCTTACGGTACCCTATATGATTTACACATTGGTCACGAGATTGGCCAATATAGGGGCAACGAACTGGATAATACATAGGACAAAAGACTAAACAAAAACATAAATTCTTGTCCTTAATTTTTTTTTTTATTTGGAATACAGTCTACTTGGTTGGAAGGTTATGTTTGACTTCAACTCAAGATTTTTTTTCTTTTTCTTGAAGAGGCTCAAGAAATATTTCTTGAGTTGAGACACGTGAAATAGGTCGCGGAGGCAATGTTAACTGATACC

Breakspear, Andrew, Chengwu Liu, Sonali Roy, Nicola Stacey, Christian Rogers, Martin Trick, Giulia Morieri, et al. 2014. “The Root Hair ‘Infectome’ of Medicago Truncatula Uncovers Changes in Cell Cycle Genes and Reveals a Requirement for Auxin Signaling in Rhizobial Infection.” *The Plant Cell* 26 (12): 4680–4701.

Gómez-Fernández, Germán O., Robin van Velzen, Jeong-Hwan Mun, Douglas R. Cook, Wouter Kohlen, and Estíbaliz Larrainzar. 2025. “Ethylene Biosynthesis in Legumes: Gene Identification and Expression during Early Symbiotic Stages.” *Journal of Experimental Botany*, February. https://doi.org/10.1093/jxb/eraf069.

Schiessl, Katharina, Jodi L. S. Lilley, Tak Lee, Ioannis Tamvakis, Wouter Kohlen, Paul C. Bailey, Aaron Thomas, et al. 2019. “NODULE INCEPTION Recruits the Lateral Root Developmental Program for Symbiotic Nodule Organogenesis in Medicago Truncatula.” *Current Biology: CB* 29 (21): 3657-3668.e5.
